# Threshold-dependent negative autoregulation of PIF4 gene expression optimizes growth and fitness in Arabidopsis

**DOI:** 10.1101/2024.12.29.630642

**Authors:** Sreya Das, Vikas Garhwal, Krishanu Mondal, Dipjyoti Das, Sreeramaiah N. Gangappa

## Abstract

PHYTOCHROME INTERACTING FACTOR 4 (PIF4) is a vital transcription factor that controls plant growth by integrating environmental signals like light and temperature. Though upstream regulators of PIF4 are known, transcriptional regulation of *PIF4* is poorly understood. Here, we demonstrate that the *PIF4* undergoes negative autoregulation. We show that *PIF4* promoter activity is more in the *pif4* mutant but significantly reduced in *PIF4* overexpression transgenic lines. Moreover, CONSTITUTIVE PHOTOMORPHOGENIC 1 (COP1), which enhances PIF4 protein stability, promotes PIF4 autoinhibition. However, Phytochrome B (phyB), a photoreceptor that decreases PIF4 stability, inhibits autoinhibition. We further develop a network-based mathematical model incorporating the PIF4 autoinhibition and other key interactions. Our modeling and data analysis reveals that PIF4 autoregulation depends on a threshold of cellular PIF4 concentration. Our model also successfully predicts the hypocotyl growth and *PIF4* promoter activity in various light and temperature conditions. Moreover, we show that the transgenic lines with enhanced PIF4 function negatively influence biomass and yield, irrespective of photoperiod and temperature. Together, the negative feedback of PIF4 dampens its own function and restraints unregulated growth. Our study thus elucidates the mechanisms of how the phyB-COP1/DET1-PIF4 module controls *PIF4* transcription in tune with the endogenous PIF4 level.

## Introduction

As sessile organisms, plants constantly monitor diurnal and seasonal changes in environments, such as light and temperature, to control their metabolism, growth, and reproduction (Franklin et al., 2014; Casal and Questa, 2018; Legris et al., 2019; Lippmann et al., 2019). Temperature and light have antagonistic relationships (Legris, 2023). Warm temperatures promote growth but compromise plant immunity and seed yield (Koester et al., 2014; Zhao et al., 2017; Burroughs et al., 2023). Therefore, balancing these two signaling pathways is critical for optimal growth and fitness (Legris et al., 2019; Casal and Fankhauser, 2023; Legris, 2023).

PHYTOCHROME INTERACTING FACTOR 4 (PIF4) is a central regulator of plant development and integrates light and temperature signals (Oh et al., 2012; Quint et al., 2016; Casal and Balasubramanian, 2019; Zhao and Bao, 2021; Quint et al., 2023). Both light and circadian clock coordinate to control *PIF4* expression by an external coincidence mechanism (Nozue et al., 2007b; Kunihiro et al., 2011). *PIF4* mRNA and protein levels are elevated in response to warmth (Koini et al., 2009; Kumar et al., 2012). PIF4 overexpression transgenic lines show constitutive thermomorphogenic response even under cooler temperatures (Lorrain et al., 2008; Kumar et al., 2012; Gangappa et al., 2017). Therefore, tight control of PIF4 levels within a range is critical for optimal growth and reproduction. Several upstream regulators, such as phytochrome B (phyB), EARLY FLOWERING 3 (ELF3), ELONGATED HYPOCOTY 5 (HY5), and CONSTITUTIVE PHOTOMORPHOGENIC 1 (COP1) / DE-ETIOLATED 1 (DET1) / SUPPRESSOR OF PHYTOCHROME A-105 (SPA) have been shown to control PIF4 activity (Casal and Balasubramanian, 2019; Delker et al., 2022). The phyB degrades PIF4 in a red-light-dependent manner(Huq and Quail, 2002a; de Lucas et al., 2008; Lorrain et al., 2008). ELF3, a component of the evening complex, inhibits *PIF4* transcription and sequesters it by forming heterodimers (Nusinow et al., 2011; Box et al., 2015; Nieto et al., 2015). Similarly, HY5 inhibits the transcriptional capability of PIF4, probably via competitive binding to target promoters (Delker et al., 2014; Toledo-Ortiz et al., 2014; Gangappa and Kumar, 2017). On the other hand, COP1/DET1/SPA promotes PIF4 protein stability (Dong et al., 2014; Gangappa and Kumar, 2017; Lee et al., 2020).

Gene regulation is critical to fine-tuning various developmental and physiological processes (Alon, 2007). Several transcription factors have evolved autoregulatory mechanisms through feedback or feedforward loops to control their function (Ohgishi et al., 2001; Rosenfeld et al., 2002; Gao and Stock, 2018). For instance, HY5, an essential activator of seedling photomorphogenesis, autoactivate to induce its transcription to amplify the signalling output (Abbas et al., 2014; Binkert et al., 2014). Contrastingly, the basic-helix-loop-helix (bHLH) transcription factor MYC2, a key regulator of growth and insect resistance pathway, negatively autoregulates its gene expression (Dombrecht et al., 2007). Interestingly, the transcription factor p53 autoregulates its gene expression both positively and negatively in various developmental contexts (Mosner et al., 1995; Wang and El-Deiry, 2006; Lu, 2010).

PIF4 can indirectly autoregulate its function by activating the gene expression of its inhibitors (Lorrain et al., 2008; Hornitschek et al., 2009; Kathare et al., 2020; Lee et al., 2021). Though upstream regulators of PIF4 are well studied, an open question is whether PIF4 can regulate its own expression and what can be the nature of such feedback. Here, by combining detailed experimental analysis with mathematical modeling, we show that PIF4 negatively autoregulates its promoter activity in a concentration-dependent manner. We examined the *PIF4* promoter activity in various genetic backgrounds by perturbing known components interacting with PIF4 signaling. Our data suggests that PIF4 inhibits its own expression when the cellular PIF4 concentration exceeds a threshold. The photoreceptor phyB promotes the *PIF4* promoter activity by reducing PIF4-dependent negative feedback. On the other hand, COP1/DET1 promotes PIF4 autoregulation. Based on these observations, we propose a minimal network model that qualitatively predicts the *PIF4* activity and hypocotyl growth in various environmental conditions and genetic backgrounds. Moreover, we found that negative autoregulation of PIF4 is critical to dampening the expression of its target genes related to growth and hormone signaling, which optimizes growth and fitness.

## Results

### PIF4 negatively autoregulates its own expression in a photoperiod-dependent manner

To investigate the role of PIF4 in regulating its own expression, we generated two transgenic lines (line #1 and line #2) containing *PIF4* promoter fused to beta-glucuronidase (GUS) reporter (*pPIF4:GUS)* (Figure S1A) in *Arabidopsis* wild-type (WT) background. The hypocotyl lengths of these *pPIF4:GUS* transgenic lines seedlings were similar to WT (Figure S1B). We also monitored GUS staining and activity of these lines under short-day (SD) and long-day (LD) photoperiods (Figure S1C-1F). The staining patterns of both lines are largely similar in SD (Figure S1C and 1D), though line #1 showed slightly more activity than line #2 in LD (Figure S1E and 1F). The *pPIF4:GUS* promoter activity peaked both during the day and at the end of the night in SD (Figure S1C and S1D), while in LD, it peaked during the daytime (Figures S1E and S1F), similar to the endogenous *PIF4* transcript accumulation, as reported earlier (Nozue et al., 2007a; Nomoto et al., 2012; Huang et al., 2016; Li et al., 2020). Also, the GUS activity was significantly higher in the LD than in the SD across different time points over a day (Figure S1G). We randomly chose the transgenic line #1 and introgressed it into *pif4-101* null mutant (Figures S1H-1I) and two overexpression lines *(PIF4-OE1* and *PIF4-OE2*)(Gangappa et al., 2017), and generated homozygous *pPIF4:GUS* lines under respective genetic backgrounds.

The overexpression lines showed that PIF4 protein accumulation is higher than WT, as revealed by immunoblotting data (Figure 1A). *PIF4-OE2* has relatively more PIF4 protein than *PIF4-OE1* (Figure 1A). Consistently, *PIF4-OE2* has longer hypocotyls than *PIF4-OE1* under SD and LD, respectively, albeit both were longer than WT (Figures S1J and S1K). Moreover, qPCR data suggests that the *PIF4-OE1* line has higher *PIF4* transcripts than *PIF4-OE2*, though *PIF4-OE1* has less PIF4 protein level than *PIF4-OE2* (Figures 1A and 1B). The GUS staining in the white light (WL)-grown WT seedlings under SD was mainly visible in the hypocotyls and the hypocotyl-root junction (Figure 1C). The *pif4-101* mutant showed a similar staining pattern, but the intensity was more prominent than WT in the cotyledons and the hypocotyl-root junction (Figure 1C). Consistent with this, quantitative GUS activity measurement of entire seedlings was two-fold higher in *pif4-101* than WT (Figure 1D), suggesting PIF4 may negatively regulate its own transcription. On the other hand, in *PIF4-OE1* and *PIF4-OE2*, the GUS staining was absent in the hypocotyls, though *PIF4-OE1* showed mild staining in the cotyledon tips (Figure 1C). Consistently, the quantitative GUS activity from seedlings was several-fold lower than WT in both overexpression lines (Figure 1D). Notably, the GUS activity was significantly reduced in *PIF4-OE2* than in *PIF4-OE1* (Figures 1C and 1D), suggesting that higher endogenous PIF4 concentration correlates with lower *PIF4* promoter activity. We further separately quantified the GUS activity from cotyledons, hypocotyls, and roots. Consistent with the *pPIF4:GUS* promoter activity in whole seedlings, the GUS activity in SD is more in the *pif4-101* mutant but less in overexpression lines than WT in all tissues (Figures 1E and S1L). Moreover, the GUS activity in *PIF4-OE2* is significantly reduced than the *PIF4-OE1* in all the tissues (Figures 1E and S1L).

**Figure 1.**
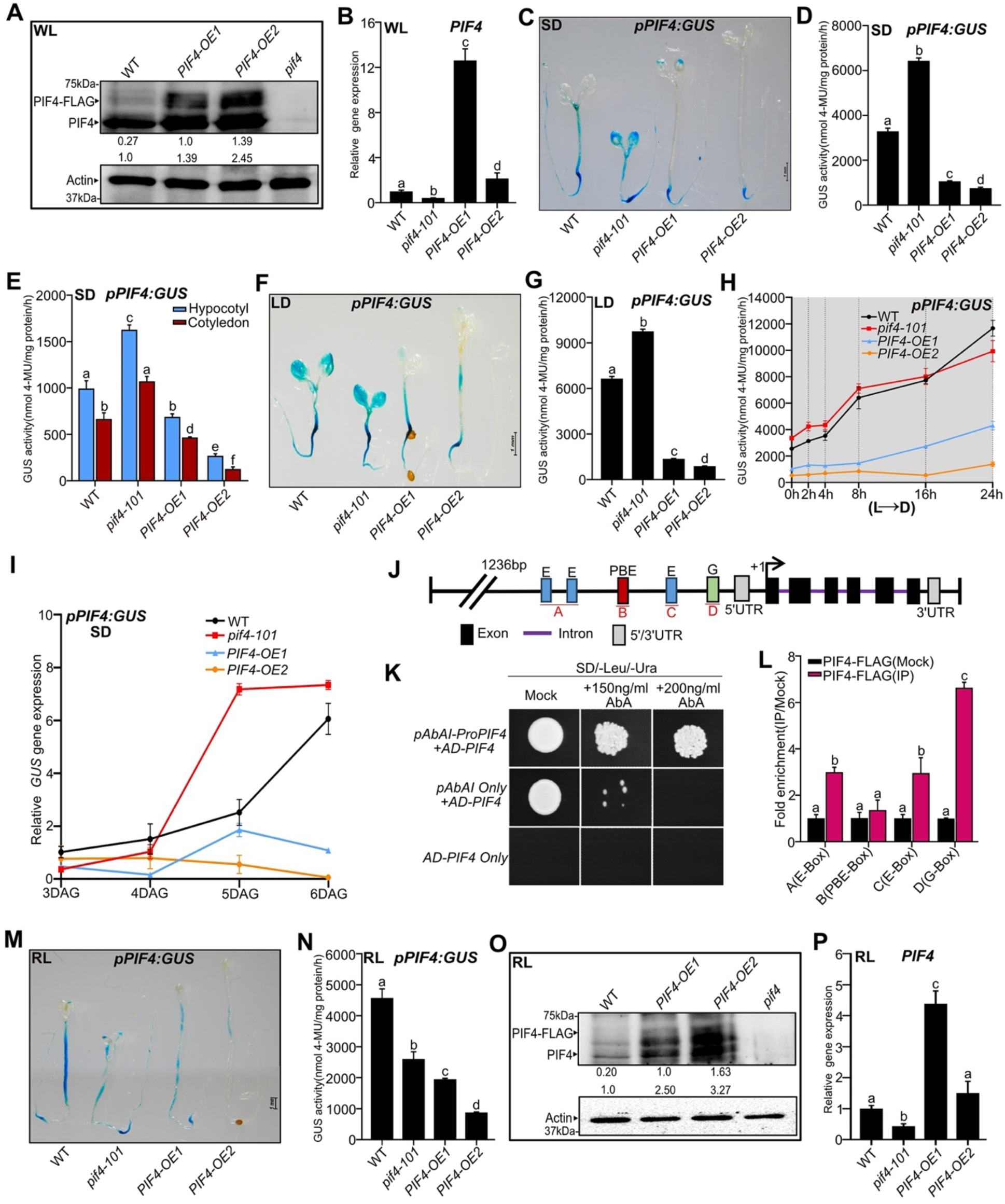
PIF4 autoinhibits its expression depending on photoperiods. (A) Immunoblot analysis of WT, *PIF4-OE1* and *PIF4-OE2* using native anti-PIF4 antibody from SD-grown seedlings at ZT23 in WL at 22°C. The endogenous PIF4 and PIF4-FLAG protein bands are indicated by arrowheads. Both the bands were considered for total PIF4 protein quantification. The total PIF4 protein level below the blots (bottom) was normalized by WT. Values for PIF4-FLAG (top) indicate the fold change compared to *PIF4-OE1*. Actin was used as a loading control. The *pif4-101* mutant was used as a negative control. (B) Relative *PIF4* transcript accumulation in six-day-old seedlings of respective genotypes grown under SD in WL and tissue harvested at ZT23. (C and D) Representative images of GUS staining (C) and quantitative GUS activity measurement (D) from six-day-old whole seedlings of WT, *pif4-101* mutant*, PIF4-OE1 and PIF4-OE2* grown under SD in WL, and seedlings were harvested at ZT23. (E) For tissue-specific GUS activity, hypocotyls and cotyledons were excised by scalpel from six-day-old whole seedlings of indicated genotypes grown under SD. Tissue was harvested at ZT23, and GUS activity was quantified. (F and G) Representative images of GUS-stained seedlings (F) and quantitative GUS activity (G) from six-day-old seedlings of indicated genotypes grown under LD in WL. (H) Genotypes were grown under constant light for five days and shifted into the dark for various durations as indicated, tissue was harvested at the specified intervals, and GUS activity was quantified. The grey zone represents the dark period. (I) The seeds were germinated for two days, and GUS transcript abundance was measured from three to six-day-old seedlings of WT*, pif4-101, PIF4-OE1* and *PIF4-OE2* grown under SD. Tissue was collected at the end of the night. (J) The schematic diagram of the *PIF4* promoter contains the E-box, PBE-box, and G-BOX. (K and L) Yeast One-Hybrid assay (K) and ChIP data (L) shows that PIF4 binds to its own promoter. ChIP was performed on six-day-old *PIF4-FLAG* seedlings grown under WL-SD (tissue was harvested at ZT23), which shows that PIF4 binds specifically to G-box and E-box (L). (M and N) Representative images of six-day-old GUS-stained seedlings (M) and GUS activity (N) in indicated seedlings grown in RL under SD. (O) Immunoblot showing endogenous PIF4 and PIF-FLAG in RL-grown seedlings under SD. Tissue was harvested at ZT23. The quantification was done as described in 1A. Actin was used as a loading control. *pif4* was used as a negative control. (P) The *PIF4* gene expression at ZT23 from six-day-old seedlings of indicated genotypes grown in RL under SD. All bar graphs represent data (mean ± SD) from three biological replicates. Different letters indicate a significant difference (one-way and two-way ANOVA with Tukey’s HSD test, P < 0.05, n ≥ 40 seedlings for GUS activity data). See also Figures S1-S3.

The *PIF4* promoter activity under LD largely followed a similar trend to SD in whole seedlings (Figures 1F and 1G) and in different tissues (Figures S1M and S1N). However, in the *PIF4-OE1*, the promoter activity in the hypocotyl was comparable to WT (Figure S1M), and in the *pif4-101* mutant, the activity in roots was comparable to WT (Figure S1N). Overall, our data suggests that increased PIF4 function enhances autoinhibition of its promoter activity irrespective of the tissue type.

Comparisons between LD and SD further suggest that PIF4 autoinhibition is regulated in a photoperiod-dependent manner (Figures S1O and S1P). As already known in the literature(Nozue et al., 2007a; Nomoto et al., 2012; Huang et al., 2016; Li et al., 2020), the PIF4 activity generally increases in the SD than in LD. Thus, according to our hypothesis of PIF4 autoinhibition, we would expect higher inhibition strength in SD than in LD. Consistent with this, we found lower GUS activity when we compared SD (ZT23) and LD (ZT4) in the WT and *pif4* mutant (Figures S1O). Moreover, at ZT4, we found increased activity in LD than in SD in both WT and *pif4* mutants (Figures S1P). However, the overexpression lines did not show marked differences in activity between SD and LD (Figures S1O and S1P), likely because the PIF4 protein level is already higher in the overexpression lines than in the WT.

Additionally, we shifted five-day-old constant light-grown seedlings to dark for different exposure times. Increased duration of dark exposure induced GUS activity in WT and *pif4-101* (Figures 1H and S1Q). In contrast, although induced by dark exposure, the promoter activity in the *PIF4-OE1* line was markedly lower than in WT (Figures 1H and S1Q). Moreover, in the *PIF4-OE2* line, the promoter activity stayed very low and showed no induction to dark treatment (Figures 1H and S1Q). Thus, our data confirms that cellular PIF4 concentration can negatively affect its promoter activity.

We also confirmed the effect of PIF4 on its autoinhibition using *pPIF4:LUC* promoter-reporter (Murcia et al., 2022). We introgressed the *pPIF4:LUC* into *pif4-101* and *PIF4-OE2* and identified double homozygous lines. Quantification of luciferase (LUC) activity as measured by luminescence from six-day-old seedlings grown in WL under SD photoperiod reveals that the *pPIF4:LUC* promoter activity was significantly upregulated in *pif4-101* but downregulated in *PIF4-OE2* (Figure S1R), further confirming PIF4 autoinhibition.

To further understand *PIF4* autoinhibition linked to protein concentration, we introduced *pPIF4:GUS* transgene into *35S:PIF4-HA* transgenic line background (Nozue et al., 2007a; Lorrain et al., 2008), which has exaggerated growth phenotypes (Nozue et al., 2007a; Lorrain et al., 2008; Gangappa and Kumar, 2018). We compared the hypocotyl phenotype of *35S:PIF4-HA* (referred to as *PIF4-OE3*) with *PIF4-OE1* and *PIF4-OE2* and found that *PIF4-OE3* has longer hypocotyls than *PIF4-OE1* and *PIF4-OE2* (Figure S2A). Moreover, qPCR data using primers specific to *5’ UTR* and part of the gene revealed that *PIF4-OE3* has lower endogenous *PIF4* transcript than *PIF4-OE1*, *PIF4-OE2* and WT (Figure S2B). Further, GUS staining in *PIF4-OE3* was hardly visible in any parts of the seedlings, similar to *PIF4-OE2* in both SD and LD (Figures S2C and S2E). Interestingly, the *pPIF4:GUS* promoter activity in *PIF4-OE3* had significantly reduced than *PIF4-OE1* and *PIF4-OE2* both in SD and LD for whole seedlings (Figures S2D and S2F) as well as in different tissues (Figures S2G-S2J).

Similar to seedlings, the *pPIF4:GUS* activity was prominent in the juvenile plants (three-week-old) of both the lines (#1 and #2) grown in WL and showed similar expression patterns and GUS activity (Figure S3A and S3B). Further, analysis of the GUS activity of leaves and stems of six-week-old adult plants showed that the *pif4-101* mutant still has significantly higher GUS activity than overexpression lines (Figures S3C-S3H), though it has lower activity than WT. This suggests that autoinhibition of *PIF4* promoter still persists in the adult stage, albeit to a lesser extent.

Further, to capture the dynamics of negative feedback activation, we quantified the GUS transcripts in the *pPIF4:GUS* line over six days (Fig 1I). We found that the GUS transcript level in the WT steadily increased from day 3 to day 6 (Fig 1I). However, the GUS transcript level in *PIF4-OE1* was significantly lower than the WT at the beginning and slightly increased on the 5th and 6th day but still stayed significantly lower than WT (Fig 1I). This data is consistent with the GUS activity measured on day 6 (Fig 1D) and suggests that autoinhibition of *PIF4* is active in *PIF4-OE1*. Moreover, the *GUS* transcript level is always lower in *PIF4-OE2* than in WT throughout the duration (Fig 1I), and it is further reduced than *PIF4-OE1* on day 5 and day 6 (Fig 1I). These results suggest that an increased PIF4 accumulation over time results in stronger autoinhibition on the fifth and sixth days.

As our results indicate *PIF4* self-inhibition, we examined if the PIF4 protein can directly associate with its own promoter. The *PIF4* promoter contains cis-acting elements, such as E-boxes, PBE-box and G-box, which could be targeted by PIF4 (Oh et al., 2012) (Figure 1J). When we transformed *pPIF4:AUR1-C* promoter-reporter constructs along with PIF4 as an effector, PIF4 could activate the *AUR1-C* gene expression and the growth of yeast in the presence of Aureobasidin A (AbA), suggesting that PIF4 could bind to its promoter (Figure 1K). Further, Chromatin Immunoprecipitation (ChIP) assay using six-day-old *pPIF4:PIF4-FLAG* transgenic seedlings and subsequent qPCR analysis revealed that PIF4 has more affinity to the G-box than the E-box but not to the PBE-box (Figure 1L).

### Effect of red light on PIF4 autoinhibition

As red light (RL) is the vital regulator of the PIF4 function(Huq and Quail, 2002b), we also analysed the GUS activity in RL-grown seedlings. We found that GUS staining and activity were reduced in the *pif4-101* mutant than in WT (Figures 1M and 1N), unlike in WL. However, the promoter activity in overexpression lines was significantly reduced than WT, similar to WL (Figures 1M and 1N). Moreover, both *PIF4-OE1* and *PIF4-OE2* have higher *PIF4* proteins than the WT, while the *PIF4* protein level is increased in *PIF4-OE2* than in *PIF4-OE1 (*Figures 1O). However, the *PIF4* transcript in *PIF4-OE1* is higher than the WT, though the transcript in *PIF4-OE2* is comparable to the WT (Figure 1P). Thus, both in WL and RL, the PIF4 protein and transcript levels do not have a one-to-one correspondence, likely due to the underlying negative feedback (Figures 1A, 1B, 1O and 1P). This might suggest that the autoinhibition of *PIF4* promoter activity depends on a threshold of PIF4 concentration. For instance, the higher protein level in *PIF4-OE2* than in *PIF4-OE1* (Figures 1A and 1O) leads to stronger inhibition of promoter activity in *PIF4-OE2* than *PIF4-OE1* (Figures 1D and 1N), hence, lower *PIF4* transcripts in *PIF4-OE2* than *PIF4-OE1* (Figures 1B and 1P). To check this hypothesis, we next developed a mathematical model of the threshold-dependent auto-regulation of *PIF4* activity.

### A mathematical model of PIF4 autoinhibition suggests that *PIF4* promoter activity is linked to the endogenous PIF4 threshold

Based on a previous study(Nieto et al., 2022), we developed a model of PIF4 autoregulation to understand its role in hypocotyl growth (Figure 2A). We assumed that PIF4 autoinhibits its expression when the cellular PIF4 concentration is above a threshold. Our model is based on a minimal genetic network of key regulators, ELF3, phyB, and COP1, which affect PIF4 activity. We incorporated known regulations on PIF4, such as (i) ELF3-dependent repression of *PIF4* transcription, (ii) phyB-dependent inhibition of PIF4 activity, and (iii) COP1-dependent stabilization of PIF4. As reported previously(Jung et al., 2016; Legris et al., 2016), we considered the photoactivated form of phyB, which becomes inactive in the dark. Our model also includes light-dependent synthesis of phyB and COP1. We modelled the ELF3 synthesis by an oscillatory function since ELF3 dynamics links the circadian clock with the diurnal cycle(Nusinow et al., 2011). We also incorporated the GUS activity, which is driven by a transgenic promoter-reporter apart from the endogenous *PIF4* promoter (Figure 2A). This transgenic promoter is also negatively regulated by PIF4, dependent on a threshold PIF4 concentration. Finally, we incorporated a variable presenting the hypocotyl growth (measured in mm) as the outcome of PIF4-dependent activation of growth genes (see the detailed mathematical description in Materials and Methods).

**Figure 2.**
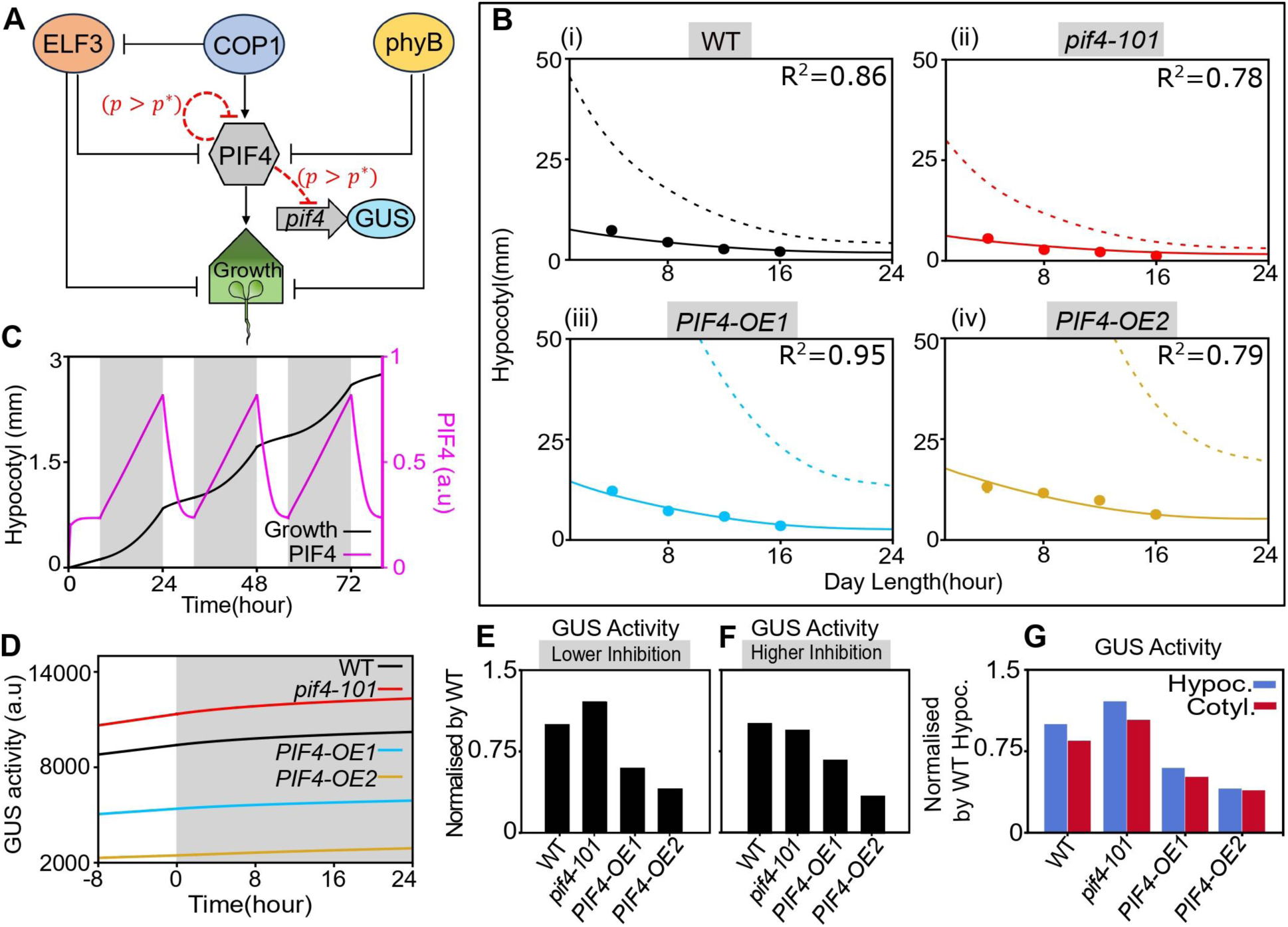
A mathematical model of PIF4 autoregulation captures the experimental observations of hypocotyl growth and promoter activity. (A) A schematic network of our model showing the regulations on PIF4. The red dashed lines denote a PIF4 autoinhibition above a threshold PIF4 concentration (see Materials and Methods for details). (B) The hypocotyl length versus day length in four genotypes in WL at 22°C. Solid lines represent predictions from our model, assuming PIF4 autoinhibition. Dashed lines denote the predictions without autoinhibition. Filled circles are experimental data for the day lengths 4,8,12, and 16 hours. (n = 20; Error bars represent Standard Deviation). The goodness of fit is shown as *R*^2^ values. (C) The hypocotyl growth and PIF4 concentration over time, predicted in our model for WT. The bright and shaded regions indicate light and dark periods within a diurnal cycle at 22°C. (D) Simulated temporal dynamics of GUS activity for four genotypes during the light-dark transition (also see Figure 1H). The grey zone represents the dark period. (E, F) Predicted GUS activity, normalized by the WT value, under SD in WL, assuming either low (E) (*P*_*self*_ = 25) or high (F) (*P*_*self*_ = 75) autoinhibition strength of PIF4 activity. (G) Predicted GUS activities in cotyledon (red) and hypocotyl (blue), normalized by the WT value in hypocotyl (compare with Figure 1E). We assumed the autoinhibition strength in cotyledon (*P*_*self*_ = 15) is lower than in hypocotyl (*P*_*self*_ = 25). Other model parameters are summarized in Tables S1 and S2.

We used our model to predict the measured hypocotyl length and GUS activity in different genetic backgrounds. The model was first fitted to wildtype data from seedlings grown under different photoperiods at 22°C to estimate WT parameters (Figure 2B(i), parameters are given in Table S1 and Table S2 in Supplemental Data). We then quantitatively predicted the hypocotyl lengths in *pif4-101* mutant and overexpression lines by varying distinct synthesis rates of PIF4 in different genotypes (Figure 2B(ii)-(iv)). Here, we assumed that the threshold concentration of PIF4, above which the autoinhibition is active, may vary with temperature and other genetic perturbations (see Table S2). Our theoretical predictions matched well with the hypocotyl data when we assumed the negative feedback on PIF4 (Figure 2B). Notably, without feedback, the predicted hypocotyl lengths are much higher in all genotypes (dashed lines in Figure 2B(i)-(iv)). This indicates that the negative feedback on PIF4 is necessary to limit the unbounded growth within a normal range. Also, our model displayed the oscillatory behaviours of PIF4 concentration and hypocotyl growth (Figure 2C) with the diurnal cycle, as reported previously(Nieto et al., 2022).

Moreover, we simulated the experimental condition where seedlings were grown at a constant WL for five days and then transferred into the dark for one day. Our model prediction is qualitatively similar to the experimental observation that overexpression lines show much lesser GUS activity than the wildtype and *pif4-101* during the long dark treatment (Figures 1H and 2D). However, an exact quantitative comparison of the GUS activity with experimental data cannot be performed since our model’s GUS activity is measured in arbitrary units.

Next, we checked the hypothesis of whether a threshold-dependent PIF4 autoinhibition could explain the differential GUS activity observed under WL and RL (Figures 1D and 1N). Note that the relative level of *PIF4* mRNA in the overexpression lines under RL is lower than the WL (Figures 1B and 1P). Thus, according to our hypothesis of PIF4 autoinhibition, we may expect stronger negative feedback in RL than in WL. Therefore, we simulated the WL and RL conditions by increasing the strength of autoinhibition in RL compared to WL. This assumption was sufficient to theoretically reproduce the observed differential GUS activity in WL and RL (Figures 2E and 2F). Moreover, our model can also explain the experimentally observed tissue-specific GUS activities in cotyledon and hypocotyl by assuming that the autoinhibition strength in cotyledon is lower than in hypocotyl (compare Figure 1E and 2G). This assumption is based on our data, which shows lower GUS staining in cotyledons than in hypocotyls (Figure 1C). Overall, our model analysis suggests that PIF4 autoinhibits its expression in a threshold-dependent manner.

### The COP1/DET1 module is essential for the *PIF4* gene expression and promotes its autoinhibition

COP1/DET1 module, a master regulator of photomorphogenesis (Deng et al., 1991; Deng et al., 1992; Pepper et al., 1994; Lau and Deng, 2012; Han et al., 2020), promotes PIF4-mediated thermosensory hypocotyl growth by stabilizing the PIF4 protein (Delker et al., 2014; Gangappa and Kumar, 2017; Park et al., 2017). To understand the role of COP1 and DET1 in regulating *PIF4* promoter activity, we introgressed *pPIF4:GUS* transgene into *cop1-4*, *cop1-6*, and *det1-1* mutants and COP1 overexpression (*35S:COP1*)(Holm et al., 2001) backgrounds. In *cop1-4* and *cop1-6* seedlings grown in SD photoperiod, the GUS staining was not visible in any tissues, compared to WT (Figure 3A). In the *det1-1* mutant, the intensity was visibly reduced, though the staining pattern was largely similar to the WT (Figure 3A). Consistently, the GUS activity decreased several-fold in the *cop1-4, cop1-6* and *det1-1* mutants (Figure 3B), implying that COP1 and DET1 positively regulate *PIF4* promoter activity. Therefore, it is expected that increased COP1 activity would increase *pPIF4:GUS* activity. However, surprisingly, the promoter activity was significantly reduced in the *35S:COP1* than in the WT (Figures 3A and 3B). Moreover, qRT-PCR data also suggested that the *PIF4* transcript level was significantly lower than WT in *cop1-4, cop1-6, det1-1* and *35S:COP1* backgrounds (Figure 3C). This unexpected result of reduced GUS activity and *PIF4* transcript in *35S:COP1* can be explained by PIF4 concentration-dependent autoinhibition. By assuming a threshold-dependent PIF4 autoinhibition, our model prediction qualitatively matched the experimental GUS activity (Figure 3D). In line with this, PIF4 stability in *35S:COP1* was higher than the WT and other mutants (Figure 3E). Like SD, the effect of COP1/DET1 on *pPIF4:GUS* activity was also broadly similar under LD-grown seedlings, as observed in experiments and our model (Figures 3F-3H). Furthermore, the phenotypic output measured by hypocotyl lengths in all genotypes was consistent with our hypothesized model of PIF4 autoinhibition since our prediction and experimental data matched well for both SD and LD (Figures 3I, S4A and S4B).

**Figure 3.**
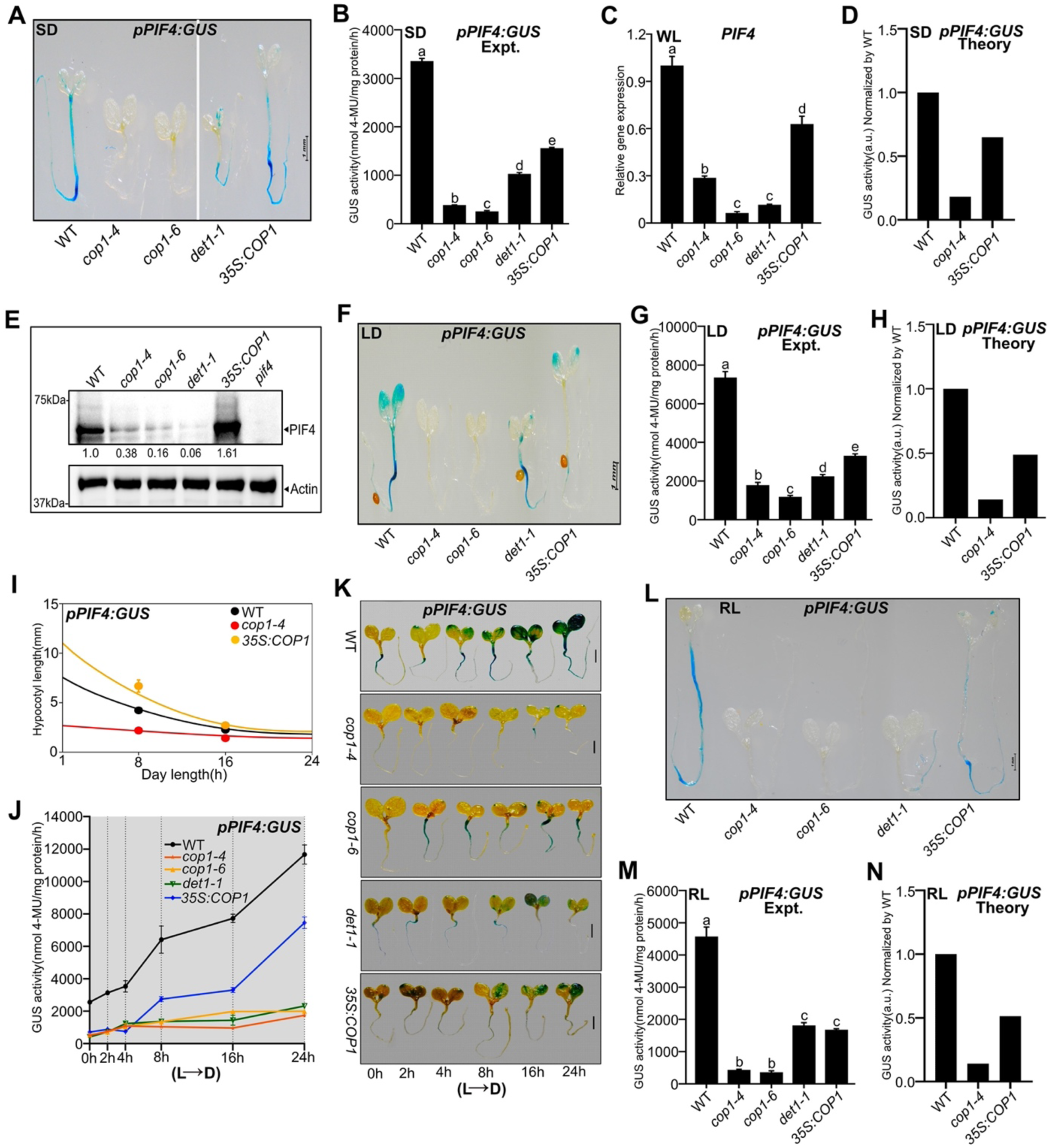
COP1/DET1 module promotes PIF4 autoinhibition. (A and B) The *pPIF4:GUS* stained images (A) and GUS activity (B) of six-day-old seedlings of indicated genotypes grown under SD in WL and seedlings were harvested at ZT23. In A, the vertical line represents two separate images. (C) *PIF4* transcript level of indicated genotypes grown under SD in WL at ZT23. (D) Predicted GUS activity from our model, normalized to WT, under SD in WL. (E) Immunoblot analysis using PIF4 native antibody showing PIF4 protein levels compared to WT. Seedlings were grown under SD in WL and harvested at ZT23. Values below the blots indicate the fold-change compared to WT. The *pif4* mutant was used as a negative control. (F-H) Images of GUS-stained seedlings (F), measured GUS activity (G) at ZT4, and predicted GUS activity from the model (H) in six-day-old seedlings grown under LD in WL. (I) Comparison between predicted and measured hypocotyl lengths of indicated genotypes. Filled circles represent experimental data (n = 20; Error bars represent standard deviations). (J and K) GUS activity (J) and GUS staining images (K) of constant light-grown seedlings (22°C) transferred to dark for varying time intervals. The grey zone in ‘J’ represents the dark period. (L-N) GUS-stained images (L), measured GUS activity data (M), and predicted GUS activity from our model (N) of six-day-old seedlings grown under SD in RL (ZT23). Data are shown as mean ± SD. Different letters above the bar charts indicate a significant difference (One-way ANOVA with Tukey’s HSD test, P < 0.05). The qRT-PCR data was obtained as fold-induction relative to WT at 22°C (Two biological replicates and three technical replicates). See also Figure S4.

Moreover, constant light-grown WT seedlings, exposed to dark for various durations, showed strong induction of its promoter activity (as shown above) but compromised in the *cop1-4, cop1-6* and *det1-1* mutants (Figures 3J and 3K). However, in the *35S:COP1* transgenic line, the promoter activity, although induced more than the mutants, is significantly less than the WT (Figures 3J and 3K). Like WL, the effect of COP1/DET1 on GUS staining was roughly similar under RL SD-grown seedlings (Figure 3L), and the measured GUS activity data also qualitatively matched with the model (Figures 3M and 3N). Furthermore, the GUS staining and activity of juvenile (Figures S4C and S4D) and adult plants (Figures S4E-S4H) revealed that COP1/DET1 positively regulates *pPIF4:GUS* activity irrespective of tissue type and developmental stages.

### PHYB positively influences *PIF4* promoter activity and impedes autoinhibition

Phytochrome B is the major inhibitor of PIF4 function as it degrades PIF4 in response to RL, leading to exaggerated growth in the *phyb* mutant(Huq and Quail, 2002a; de Lucas et al., 2008; Lorrain et al., 2008; Leivar and Quail, 2011; Jung et al., 2016; Legris et al., 2016; Park et al., 2018). To understand the role of phyB on *PIF4* promoter activity, we introgressed *pPIF4:GUS* transgene into the *phyb-9* mutant and *35S:PHYB* overexpression lines. Data revealed that GUS staining and activity under SD at 22°C were significantly reduced in the *phyb-9* mutant but were significantly enhanced in the *35S:PHYB* (Figure 4A and 4B). Our mathematical model consistently predicted this experimental trend by assuming PIF4 autoinhibition (Figure 4C). In line with this, PIF4 protein was more stable in *phyb-9* but less stable in *35S:PHYB,* than WT (Figure 4D). However, *PIF4* transcripts were higher both in *phyb-9* and *35S:PHYB* than in WT (Figure 4E). Thus, there is no linear one-to-one correspondence between the protein and transcript levels, again indicating underlying negative feedback. In *35S:PHYB,* there is much weaker negative feedback, resulting in higher *PIF4* promoter activity and likely producing more transcripts than the WT (Figure 4E). Meanwhile, in *phyb-9*, there is stronger negative feedback, leading to lower promoter activity, though producing mildly higher or comparable transcripts compared to the WT (Figure 4E).

**Figure 4.**
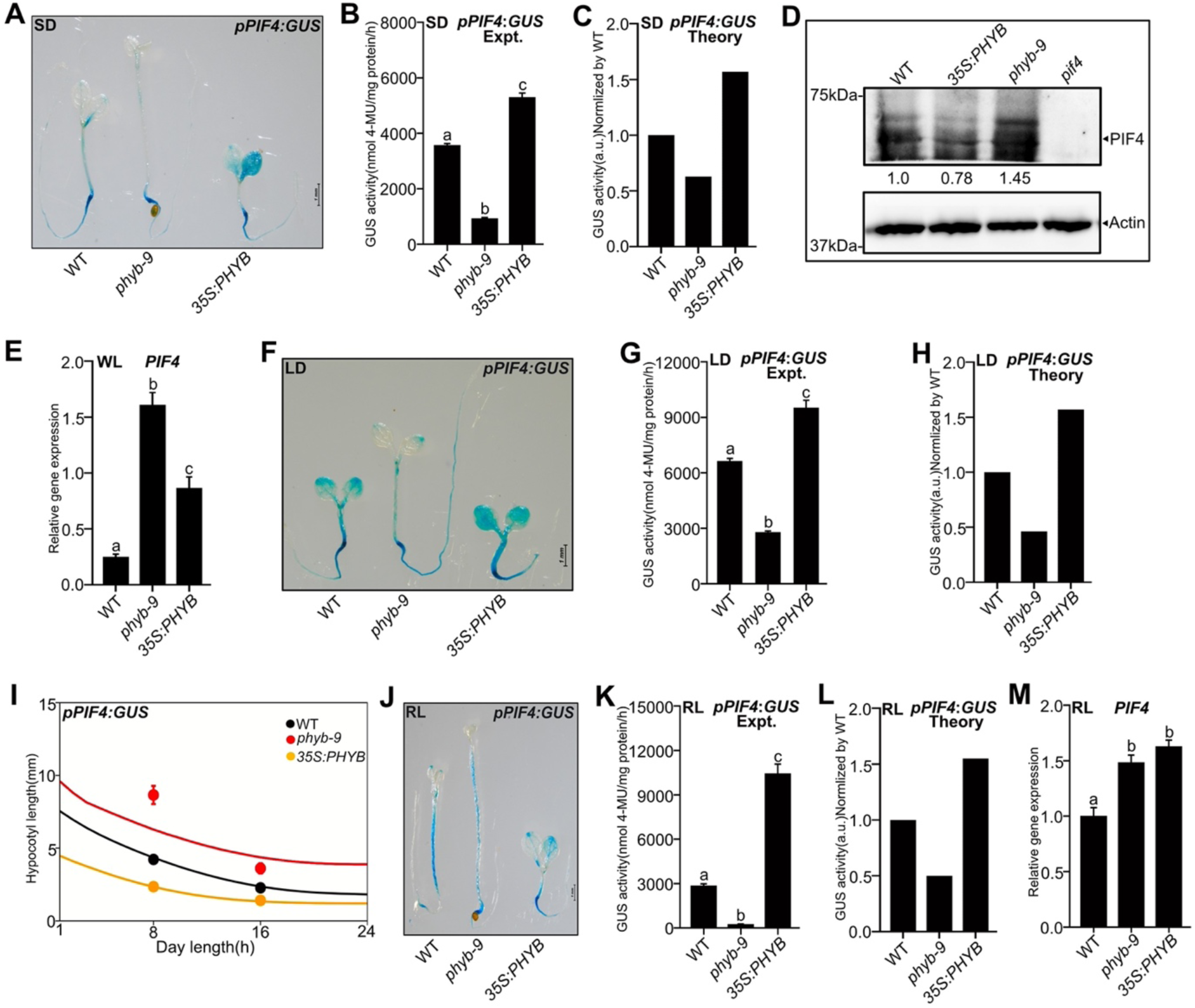
PHYB negatively regulates PIF4 autoinhibition. (A-C) Representative images of GUS staining (A), measured GUS activity (B) and predicted GUS activity from the model (C) of six-day-old WT, *phyb-9*, and *35S:PHYB* seedlings grown under SD in WL. Both for GUS staining and activity measuremnts, seedlings were harvested at ZT23. (D) The PIF4 protein levels at ZT23 as revealed by immunoblot (using anti-PIF4 antibody) in the respective genotypes grown under SD in WL. Values below the blots indicate fold-change compared to WT. Actin was used as the loading control. The *pif4* mutant was used as a negative control. (E) Real-time qPCR analysis of *PIF4* gene expression (at ZT23) from six-day-old WT, *phyb-9*, and *35S:PHYB* seedlings grown under SD in WL. (F-H) The *pPIF4:GUS* staining images (F), GUS activity measurement (G), and predicted GUS activity (H) from six-day-old seedlings grown under LD in WL. For GUS staining and activity measuremnts, seedlings were harvested at ZT4. (I) Predicted hypocotyl lengths for respective genotypes from the model. Filled circles denote experimental data (n = 20; Error bars represent standard deviations). (J-L) The representative GUS staining images (J), quantitative GUS activity (K) at ZT23 and the model predicted GUS activity (L) of six-day-old seedlings grown under SD in RL. (M) Fold change of *PIF4* transcript compared to WT under SD in RL (two biological and three technical replicates). Different letters in bar graphs indicate a significant difference (One-way ANOVA with Tukey’s HSD test, P < 0.05). See also Figure S4.

Similar to the SD data, the GUS activity under LD was also lower in *phyb-9* but enhanced in *35S:PHYB,* compared to WT (Figures 4F and 4G). This trend also matched our model prediction (Figure 4H). Consistent with the *PIF4* promoter activity, our model predicted that the hypocotyl growth should be enhanced in *phyb-9* but reduced in *35S:PHYB,* compared to WT, for both SD and LD. This prediction was consistent with the measured hypocotyl length (Figures 4I, S4I and S4J). Like the SD data in WL, the predicted and measured *pPIF4:GUS* activity was significantly reduced in *phyb*-9 but increased in *35S:PHYB* under RL (Figures 4J-4L). Also, similar to the WL, *PIF4* transcript in RL was more in both *phyb-9* and *35S:PHYB* than WT (Figure 4M), suggesting that *PIF4* autoinhibition may crucially depend on endogenous PIF4 concentration in each genotype. However, the GUS staining and activity of juvenile (Figures S4K and S4L) and adult plants (Figures S4M-S4P) showed lesser *PIF4* activity in *35S:PHYB* than WT, suggesting that phyB may have a diminished role in regulating *PIF4* activity beyond the seedling stage.

### PIF4 differentially regulates *PIF4* promoter activity in a temperature-dependent manner

PIF4 is vital for thermosensory growth and reproductive transition in Arabidopsis (Koini et al., 2009; Kumar et al., 2012). Since *PIF4* gene expression is proportional to temperature (Koini et al., 2009; Kumar et al., 2012; Delker et al., 2014; Gangappa and Kumar, 2017), we compared the *pPIF4:GUS* staining and activities in WT, *pif4-101, PIF4-OE1* and *PIF4-OE2* seedlings grown at 22°C and 27°C under SD. As expected, the *PIF4* promoter activity was higher at 27°C compared to 22°C in the WT (Figures 5A and 5B). In the *pif4-10*1 mutant, the *pPIF4:GUS* activity was significantly higher than WT at 22°C, while it was reduced at 27°C (Figures 5A and 5B). In overexpression lines, the *PIF4* promoter activity was significantly lower than WT under both 22°C and 27°C, indicating the PIF4 autoinhibition (Figures 5A and 5B). Moreover, the *PIF4* promoter activity in *PIF4-OE1* and *PIF4-OE2* lines is slightly higher at 27°C than at 22°C. These observations could be due to higher PIF4 stability at 27°C than 22°C and correspondingly increased PIF4 concentration threshold for autoinhibition. By including this hypothesis, our model could qualitatively reproduce the experimental trend of temperature dependence on *PIF4* promoter activity under SD (Figure 5C). Moreover, this hypothesis is consistent with the comparative PIF4 protein levels observed at 22°C and 27°C (Figure 5D). The tissue-specific GUS activity also showed a similar trend to the whole seedlings (Figures 5E, 5F, and S5A). Further, the activity under LD photoperiod showed a similar trend of temperature dependence in *pif4-101* and overexpression lines compared to the WT (Figures S5B and S5C). This trend was also reproduced in our model (Figure S5D). Next, we compared the GUS activities in hypocotyls, cotyledons, and roots. Similar to the whole seedling data, *pif4-101* showed reduced activity than WT at 27°C, and both overexpression lines showed a much stronger reduction in the GUS activity in all the tissues (Figures S5E-S5G).

**Figure 5.**
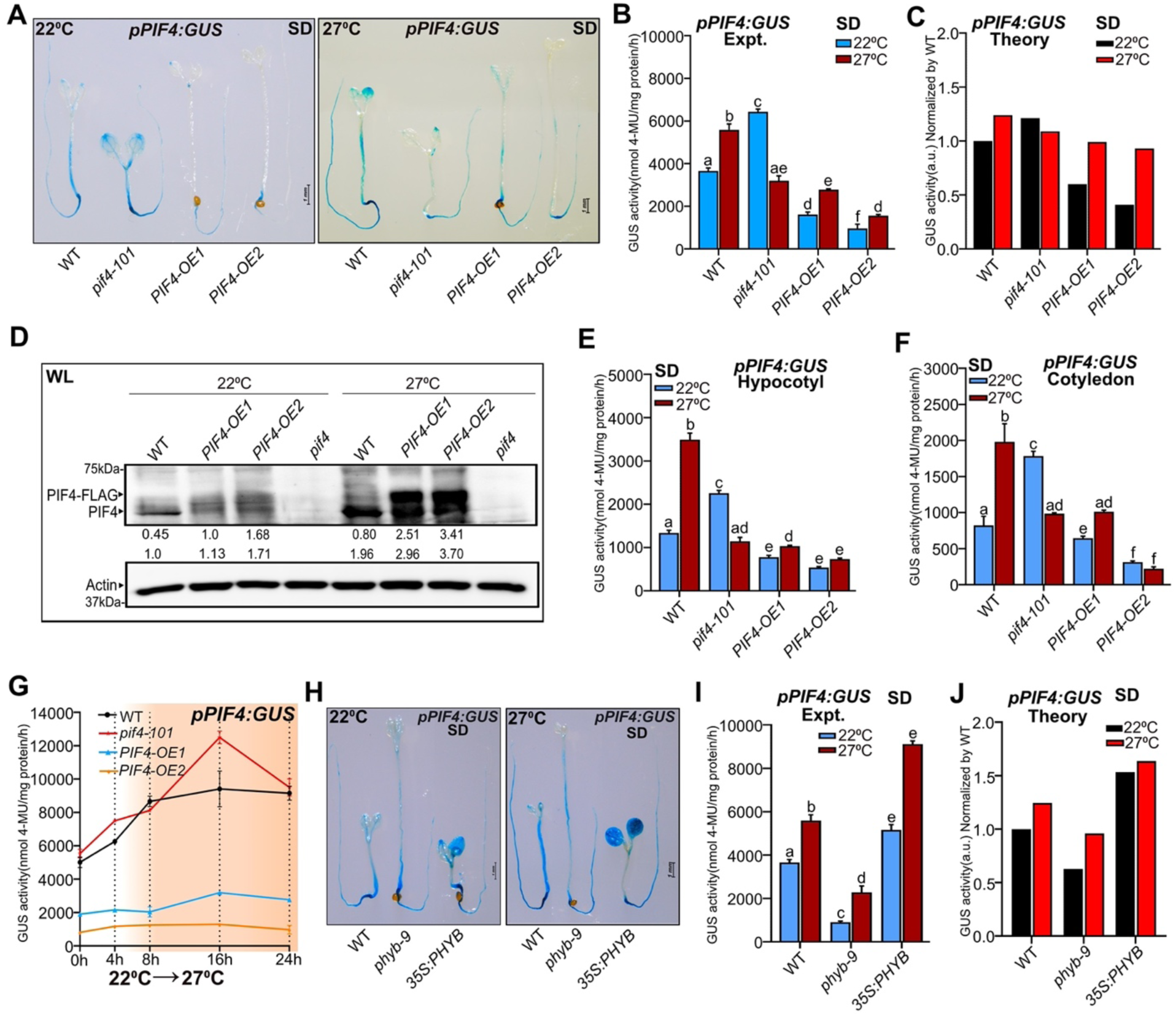
PIF4 protein level regulates *PIF4* promoter activity in a temperature-dependent manner. (A and B) Representative images of GUS-stained seedlings (A) and measured GUS activity (B) from six-day-old seedlings of respective genotypes grown under SD in WL at 22°C and 27°C, and the seedlings were harvested at ZT23. (C) Predicted GUS activity from the model at 22°C and 27°C for SD-grown seedlings. (D) PIF4 protein level from immunoblot (anti-PIF4 antibody) at ZT23 in indicated genotypes of six-day-old seedlings grown at 22°C and 27°C under SD. The endogenous PIF4 and PIF4-FLAG bands are indicated by the arrow. Values below the blots indicate fold-change compared to WT (22°C) to calculate endogenous PIF4 protein level (bottom). The values for the PIF4-FLAG band (top) were shown underneath each immunoblot, and normalization was done by setting *PIF4-OE1 (*22°C) as 1. Actin was used as the loading control. *pif4* mutant was used as a negative control. (E and F) Tissue-specific quantitative GUS measurement from six-day-old seedlings grown under SD at 22°C and 27°C at ZT23. The whole seedlings were cut into two sections, i.e. hypocotyl (E) and cotyledon (F), and GUS activity was carried out individually (see Materials and Methods). (G) GUS activity of seedlings, grown at 22°C for five days and shifted to 27°C for various time points. The yellow-shaded region represents warmer temperatures. (H-J) Representative GUS stained images (H), measured GUS activity (I) at ZT23, and model-predicted GUS activity (J) of WT, *phyb-9*, and *35S:PHYB* seedlings grown at 22°C and 27°C in WL under SD. Different letters in bar charts indicate a significant difference (two-way ANOVA with Tukey’s HSD test, P < 0.05). See also Figure S5.

Likewise, the temperature-dependent regulation of *PIF4* promoter activity became clearer when we grew the seedlings at 22°C for six days in SD and exposed them to 27°C for different time duration. Here, the *PIF4* promoter was induced over time in *pif4-101,* similar to the WT (Figure 5G and S5H). However, in overexpression lines, the promoter activity stayed several folds lower than WT (Figure 5G and S5H).

Since warmer temperature-mediated inhibition of phyB activity results in enhanced PIF4 activity(Jung et al., 2016; Legris et al., 2016), we checked *pPIF4:GUS* activity in *phyb-9* and *35S:PHYB* genotypes under both SD and LD. In the *phyb-9* mutant, the *pPIF4:GUS* activity was lower than WT at 22°C and 27°C (Figures 5H, 5I, S6I, and S6J), similar to PIF4 overexpression lines (Figures 5K and 5L). On the other hand, the *pPIF4:GUS* activity in *35S:PHYB* is higher than WT at 22°C and 27°C (Figures 5H, 5I, S6I, and S6J), again suggesting that higher phyB level leads to reduced autoinhibition. We further checked if our model could reproduce this experimental trend by assuming less phyB-mediated inhibition of PIF4 at higher temperatures. Our model successfully captured the effect of phyB on PIF4 autoinhibition in both temperatures under SD and LD (Figures 5J and S5K).

### PIF4 protein dynamics reveal differential autoinhibition of its promoter activity during day and night

Since PIF4 protein accumulation can vary over time and with temperatures, we monitored the PIF4 dynamics in the WT and overexpression lines (*PIF4-OE2*) at 22°C and 27°C under SD conditions. Our immunoblot analysis shows that the PIF4 protein accumulation peaks during the day in WT (around ZT2 at 27°C and ZT4 at 22°C), while the peaks appear slightly later in the overexpression lines (around ZT4 at 27°C and ZT8 at 22°C) (Figure 6A). In all cases, the PIF4 protein level drops slightly after the peak and is maintained throughout the night (Figure 6B and 6C). Also, as expected, the overall PIF4 protein level in *PIF4-OE2* is higher than the WT throughout the diurnal cycle. Moreover, *PIF4-OE2* shows a slightly higher peak in PIF4 protein compared to the WT. We also measured the *pPIF4:GUS* activity at the same time points at 22°C and 27°C under SD conditions. In the WT, the promoter activity peaks around ZT4 at 27°C and ZT8 at 22°C as reported (Li et al., 2024), which comes slightly later than the peak in PIF4 protein accumulation (Figures 6D and 6E). This suggests that the accumulation of proteins beyond a threshold dampens the promoter activity at the respective time points. Moreover, the promoter activity in *PIF4-OE2* is much lower than in WT throughout the day and night, suggesting strong autoinhibition since relative protein accumulation in these lines is maintained higher than in WT.

**Figure 6.**
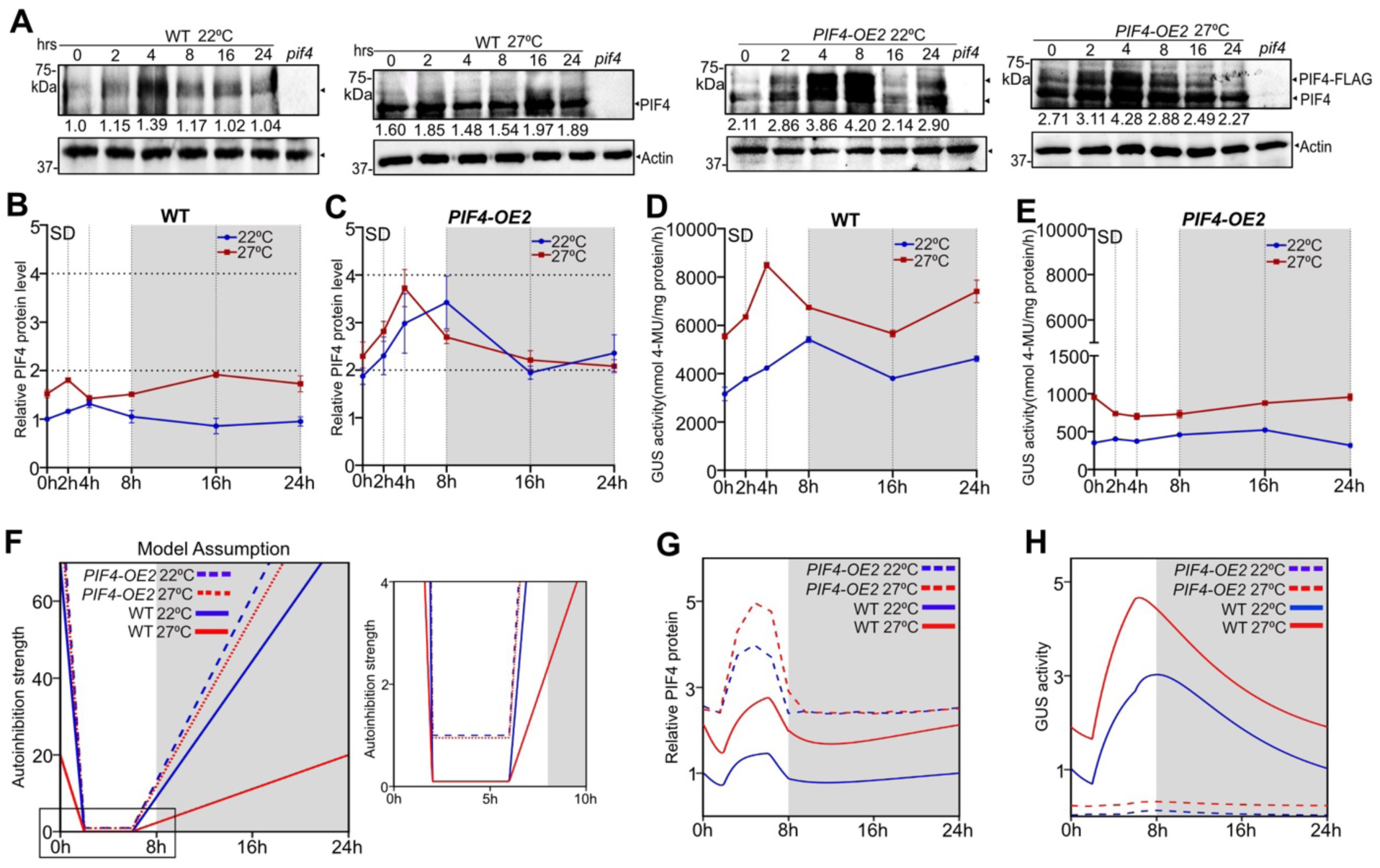
Dynamics of PIF4 protein accumulation and autoinhibition in response to temperature. (A) Immunoblot analysis at various time points of the PIF4 protein levels from seedlings of WT and *PIF4-OE2* grown at 22°C (up) and 27°C (down) under SD in WL. Tissue was harvested at ZT0 on the sixth day and grown for 24 hours. Arrowheads indicate PIF4 and PIF4-FLAG bands, which were considered for total PIF4 protein quantification. Values below the blots indicate fold-change compared to WT at 22°C, at ZT0. Actin was used as a loading control, and *pif4* mutant as a negative control. All the immunoblotting experiment was performed at teh same time and maintained the identical conditions thrpughout the procedure. (B and C) The dynamics of relative protein levels of WT (B) and *PIF4-OE2* (C) at 22°C and 27°C. The relative protein levels were normalized by the WT value at 22°C, at ZT0 (data represents mean ± SEM for 2 biological replicates). The dark period is shown in grey. (D and E) The measured GUS activities of WT (D) and *PIF4-OE2* (E) grown at 22°C and 27°C under the same conditions as described in 6.A. Data represents mean ± SD for n ≥ 40 seedlings. (F) The model assumption shows that the PIF4 autoinhibition strength varies over time. The autoinhibition strength is assumed to be higher during the night and at the beginning of the day compared to the mid-day (left). Overall, the strength is higher in *PIF4-OE2* than in the WT. The autoinhibition strength at 27°C for WT is also assumed to be lesser than at 22°C, but this difference is negligible for *PIF4-OE2* (right). See Table S3 for parameter values.(G and H) Model predictions for PIF4 protein (G) and the GUS activity (H) over time at 22°C and 27°C. Data are normalized to respective WT values at 22°C, at ZT0. Grey shade denotes the dark period.

Since PIF4 protein accumulation and corresponding promoter activity are dynamic, the autoinhibition strength may vary over time. The promoter activity in the WT drops in the middle of the night and the beginning of the day, suggesting that autoinhibition could be higher in these time points as compared to the mid-day (Figure 6D). Moreover, the promoter activity in the WT at 27°C is higher than at 22°C, suggesting lower autoinhibition strength at higher temperatures (Figure 6D). We thus assumed a similar variation of PIF4 autoinhibition strength in our mathematical model (Figure 6F). With this assumption, our model can qualitatively predict the similar dynamics of PIF4 protein and GUS activity for both WT and *PIF4-OE2* at 22°C and 27°C (Figures 6G and 6H). Together, our data reveal the complex interplay of PIF4 protein accumulation and the strength of negative feedback varying over time and with temperature.

### Elevated PIF4 dampens the expression of growth-promoting genes

PIF4-mediated promotion of thermosensory growth is linked to the direct activation of many genes involved in hormone biosynthesis, signalling, cell-wall elongation and floral transition(Koini et al., 2009; Franklin et al., 2011; Kumar et al., 2012). We checked how increased PIF4 at high temperatures affects the expression of some key targets of PIF4, such as *YUCCA8* (*YUC8*) and *INDOLE-3-ACETIC ACID INDUCIBLE 29* (*IAA29*). In WT, *YUC8* and *IAA29* expression was significantly upregulated at 27°C than 22°C (Figures 7A and 7B). However, compared to the WT, their expression in the *pif4*-101 mutant was significantly lower at both 22°C and 27°C (Figures 7A and 7B). Therefore, PIF4 is essential to activate these growth-promoting genes. In PIF4 overexpression lines, *YUC8* and *IAA29* showed significant upregulation than WT, irrespective of temperatures (Figures 7A and 7B). Notably, their expression was significantly reduced at 27°C than 22°C, suggesting that elevated PIF4 stability dampens the expression of these genes involved in growth.

**Figure 7.**
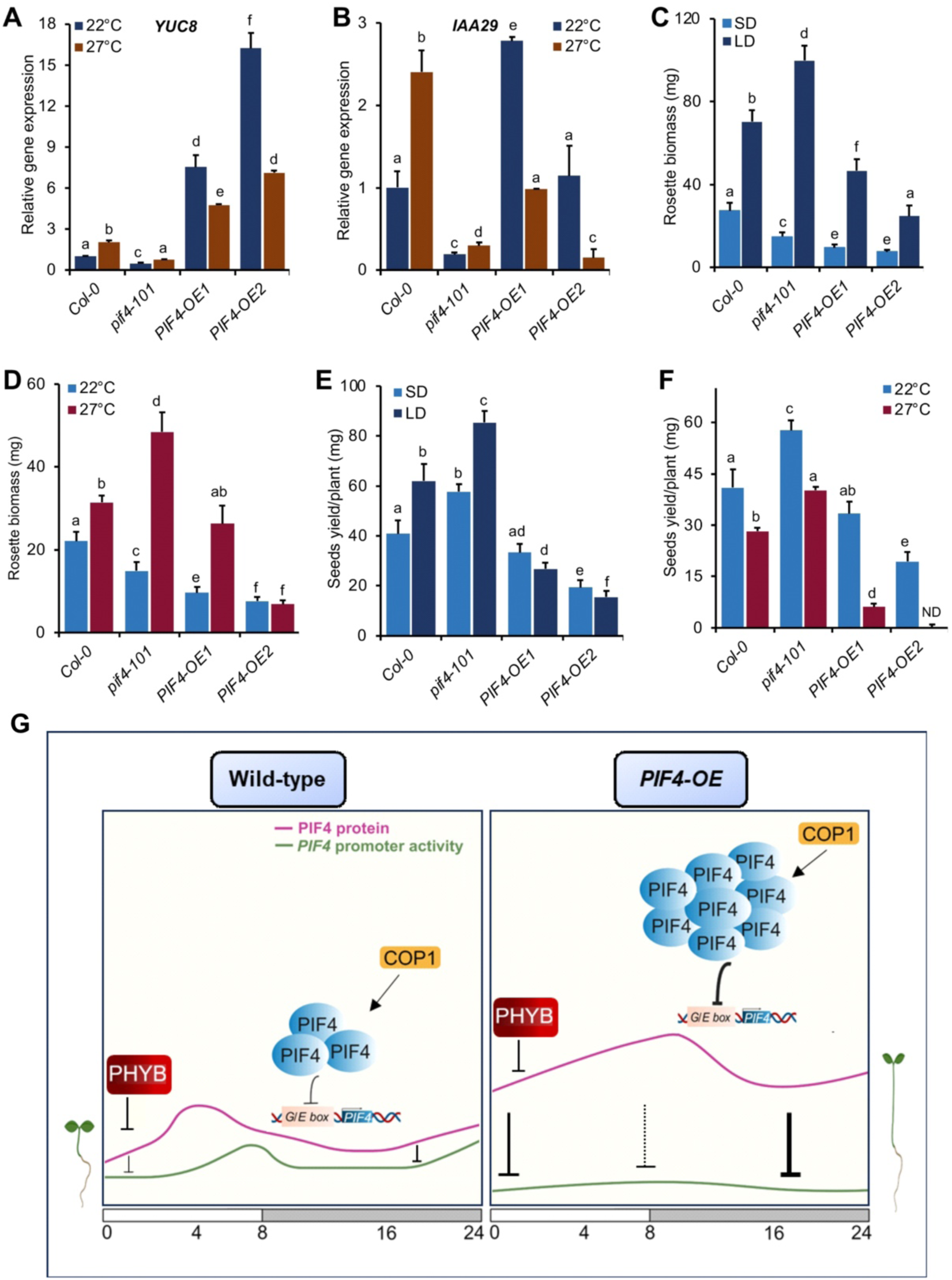
Differential expression of growth-promoting and growth-inhibiting genes optimizes biomass production and seed yield. (A and B) Expression of *YUC8* (A) and *IAA29* (B) genes at ZT23 as measured by qPCR in respective genotypes grown under SD in WL at 22°C and 27°C. *YUC8* and *IAA29* are temperature-induced and growth-responsive genes. (C and D) Rosette biomass of WT, *pif4-101, PIF4-OE1* and *PIF4-OE2* lines grown under different photoperiods at 22°C (C) and at different temperatures under SD (D). (E and F) Seed yield of indicated genotypes grown in different photoperiods at 22°C (E) and at different temperatures under SD (F). Data represent mean±SD (n≥10 plants). Different letters in bar charts indicate a significant difference (two-way ANOVA with Tukey’s HSD test, P < 0.05). (G) The graphical summary shows the dynamic nature of PIF4 protein accumulation and the corresponding autoinhibition of its promoter activity in WT and PIF4 overexpression backgrounds. In the WT, mild autoinhibition in the daytime and stronger inhibition at night. In the overexpression background, autoinhibition is stronger overall than WT.

### PIF4 threshold-dependent autoinhibition is key for optimal biomass and seed yield under varying photoperiod and temperature

Warm ambient temperature accelerates growth and development while compromising seed set and yield(Peng et al., 2004; Zhao et al., 2017). Though PIF4 is critical for warm temperature growth, its role in regulating biomass and seed yield remains elusive. To investigate this, we measured the rosette biomass (fresh weight) of three-week-old WT, *pif4-101* and *PIF4-OE* lines under varying photoperiods and temperatures. We found that rosette biomass was significantly reduced in the *pif4-101* than WT under SD, while it accumulated more biomass than WT under LD (Figure 7C). However, increased PIF4 activity in *PIF4-OE1* and *PIF4-OE2* significantly reduced biomass compared to WT, independent of photoperiods (Figure 7C). Notably, the rosette biomass of *PIF4-OE2* was reduced considerably than *PIF4-OE1* under LD (Figure 7C). Also, the *pif4-101* mutant accumulated more biomass under 27°C SD than 22°C (Figure 7D). At 27°C, the biomass in *PIF4-OE1* was similar to WT, while it was significantly reduced in *PIF4-OE2* than *PIF4-OE1* (Figure 7D). Thus, the biomass depends on the concentration-dependent PIF4 autoinhibition in response to light and temperature.

Similar to biomass accumulation, we also found that PIF4 is essential for optimal seed production. The WT produced significantly more seed yield under LD than SD, and *pif4-101* produced more seeds than WT under both SD and LD (Figure 7E). Conversely, overexpression lines produced significantly fewer seeds than WT, irrespective of the photoperiod (Figure 7E). Under warm temperatures, WT plants produced significantly fewer seeds than 22°C, while *pif4-101* produced more seeds than the WT at both 22°C and 27°C (Figure 7F). Interestingly, *PIF4-OE1* produced fewer seeds at 27°C than at 22°C, while *PIF4-OE2* did not produce any seeds specifically at 27°C (Figure 7F). This suggests that increased PIF4 levels, especially under warm temperatures, are detrimental to the optimal seed set.

## Discussion

PIF4, the key integrator of multiple environmental cues such as light and temperature, is strictly regulated to control the signalling output in tune with the endogenous hormones/metabolic fluxes (Oh et al., 2012; Casal and Balasubramanian, 2019; Delker et al., 2022). Here, we identify and demonstrate a novel mechanism through which PIF4 negatively autoregulates its promoter activity to optimise growth and grain yield in response to varying photoperiods and temperatures. Consistently, the *pif4* mutant showed higher promoter activity than the WT, while the PIF4 overexpression lines displayed strongly reduced activity under both white and monochromatic red light (Figure 1A-D, 1M-P). Moreover, overexpression of phyB, which inhibits PIF4 stability, phenocopied *pif4* mutant, showing higher promoter activity (Figure 4A-H). Similarly, *phyb-9* mutant behaves like the PIF4 overexpression lines in repressing *PIF4* promoter activity (Figures 4A-4H). On the other hand, the COP1 overexpression line also showed lower promoter activity than the WT, probably due to higher *PIF4* levels enforcing autoinhibition (Figure 3A-H). Together, these data provide evidence that PIF4 autoinhibits its transcription.

A crucial insight from our study is that we cannot always expect a linear one-to-one correspondence between the PIF4 protein and its transcript levels in the presence of negative feedback. For instance, *35S:PHYB* overexpression has reduced PIF4 protein levels and hence weaker autoinhibition, leading to higher promoter activities and higher transcripts than WT. On the other hand, *PIF4-OE2* has significantly more PIF4 protein and stronger autoinhibition, resulting in lower promoter activities and lower or comparable transcripts compared to WT. However, when the PIF4 protein levels are mildly higher than WT, as in the cases of *PIF4-OE1* and *phyb-9* mutant, the correspondence between the PIF4 proteins and transcript becomes nonlinear due to negative feedback. Although we may expect lower transcripts due to higher PIF4 stability, a mild increase in PIF4 may not be enough to cross the threshold needed to lower the transcripts significantly. Also, since the autoinhibition is a dynamic process, the transcript levels for *PIF4-OE1* and *phyb-9* mutant may become temporarily higher than or comparable to the WT, highlighting the complexity of such regulation.

Notably, we observed differential promoter activity in two PIF4 overexpression lines with distinct PIF4 protein levels, suggesting that PIF4 autoinhibition depends on endogenous PIF4 concentration (Figures 1A-1G). Furthermore, *cop1* mutants showed negligible *PIF4* promoter activity compared to WT, suggesting that endogenous PIF4 level in these genotypes is very low (Figure 3A-3H). Similarly, the *pif4* mutant at higher temperatures showed reduced *PIF4* promoter activity than WT, though the activity at lower temperatures is much higher than WT (Figures 5A-5C, S5B-S5D). This differential temperature response may indicate that *PIF4* autoinhibition takes place when PIF4 concentration reaches above a threshold. In line with this, a recent report suggests that PIF4 may even autoactivate below the threshold concentration (Zhai et al., 2020). In general, the PIF4 threshold levels in cells may change depending on light and temperature, which further dictates whether the *PIF4* transcription should be activated or repressed.

We further developed a minimal network model using the hypothesis of a concentration threshold-dependent PIF4 autoinhibition, including the main interactions between PIF4, COP1, ELF3, and PHYB (Figure 2A). This model successfully predicted the measured hypocotyl lengths in various genetic backgrounds and also recapitulated the observed trends of *PIF4* promoter activity. This further supports our claim that PIF4 inhibits itself in a concentration-dependent manner. Interestingly, our model showed unbounded high hypocotyl growth without PIF4 autoregulation, emphasizing its critical role in controlling plant growth (Figure 2B). We also explored how this mechanism relates to the expression of PIF4 targets affecting growth and overall fitness, such as rosette biomass and seed yield in varying photoperiods and temperatures (Figures 7A-7F).

In summary, higher endogenous PIF4 levels promote lesser *PIF4* promoter activity and correlate with longer hypocotyl (Figure 7G). Moreover, the PIF4 protein accumulation and subsequent autoinhibition of its promoter activity are complex and dynamic (Figure 6). In WT, our data and model suggest that the autoinhibition strength is higher at night, during which PIF4 protein accumulates and passes a threshold to inhibit its own transcription. This negative feedback leads to asynchronous oscillation of PIF4 protein and corresponding promoter activity for effective maintenance of the PIF4 protein level during the night (Figure 7G). However, in overexpression lines, the relative PIF4 protein is much higher than WT, resulting in a stronger autoinhibition. Together, we propose that PIF4 undergoes autoinhibition, either regulating its own transcription in a concentration-dependent manner or via promoting downstream regulators that inhibit PIF4.

## Materials and methods

### Plant materials and growth condition

Unless otherwise specified, all genetic materials used in this study are in Col-0 background. The various mutants and transgenic lines used in the study are listed in the Key Resources Table. For various experiments in this study, seeds were surface-sterilized (70% ethanol + 0.05% Triton X-100) and germinated on Murashige and Skoog (MS; pH: 5.7) plates containing 1% sucrose (Sigma-Aldrich) and 0.8% w/v agar (Himedia Laboratories, India) following stratification for three days at 4°C in the dark. Next, seeds were transferred to 22°C under LD photoperiod (LD; 16h light/8 h dark) in the plant growth chamber for germination. Upon germination, seedlings were either transferred to 27°C or retained at 22°C for six days unless otherwise specified. The light intensity of 100 µmol m^−2^ s^−1^ was used to grow seedlings for all the experiments. Experiments were performed under dark or short-day (SD; 8h light/16 h dark), long-day (LD; 16h light/8 h dark), or constant light (24 h light) conditions as specified. For monochromatic light experiments, red light (RL) was used at 60 µmol m^−2^ s^−1^ intensity. Experiments related to monochromatic lights were performed under short-day conditions.

### Vector Construction and generation of transgenic lines

The promoter-reporter line *pPIF4:GUS* was constructed by gateway-based cloning. Genomic DNA from Col-0 was used to amplify the *PIF4* (1236 bp) promoter and the PCR fragment was cloned into pENTR/D-TOPO and inserted into the pGWB633 (carboxy-terminal GUS) vector. All recombinant clones were validated through restriction analysis and were sequence-confirmed. Then, the specific constructs were transformed into *Agrobacterium tumefaciens GV3101*. Transgenic plants were obtained by transforming Col-0 by the floral-dip method. Transgenics were selected on MS+BASTA (7.5 µg/µl) antibiotic. Two independent homozygous transgenic lines (Lines #1 and #2) were analysed for GUS activity, and transgenic line #1 was used for further analysis.

### Generation of *pPIF4:GUS* in various mutant and overexpression lines

For better understanding, we labelled PIF4-overexpressor lines and categorised them based on their hypocotyl length; *pPIF4:PIF4-FLAG-OE1 (PIF4-OE1)* has moderately long hypocotyl phenotype while *pPIF4:PIF4-FLAG-OE2 (PIF4-OE2)* has very long hypocotyls and *35S:PIF4-HA (PIF4-OE3)* has significantly long hypocotyls. For generating *pPIF4:GUS* promoter-reporter transgene in various genotypic backgrounds, we crossed *pPIF4:GUS transgenic line* separately with *pif4-101*, *PIF4-OE1, PIF4-OE2, PIF4-OE3*, *cop1-4, cop1-6, det1-1, 35S:COP1-OE, phyb-9, 35S:PHYB-GFP*, *hy5-215* and *elf3-4* to obtain F_1_ seeds. F_1_ seeds were further propagated to get F_2_. Next, F_2_ seeds were screened by plating on MS along with appropriate antibiotic selection. Desired seedling combinations were selected based on antibiotic selection and hypocotyl phenotype. The selected plants were further genotype and/or phenotype confirmed in the adult stage, along with GUS staining for the *pPIF4:GUS* transgene (Please refer to **Table S4** for primers used for genotyping). For generating *pPIF4:LUC* promoter-reporter transgene in the background of *pif4-101* and *PIF4-OE2*, we crossed with the lines separately, and F_2_ screening was performed with appropriate antibiotic selection and proceeded for homozygous lines. The putative homozygous lines were further confirmed in the next generation by antibiotic selection and phenotypic analysis (hypocotyl measurement) before further analysis.

### Hypocotyl measurements

Six-day-old seedlings of various genotypes grown under either SD or LD were used to measure hypocotyl lengths. On the sixth day, ∼20-25 seedlings per genotype were aligned on an Agar plate containing charcoal before being photographed along with scale. Later, hypocotyl lengths were measured using NIH ImageJ software (https://imagej.nih.gov/ij).

### GUS histochemical staining assay

For the GUS histochemical assay, six-day-old seedlings, three-week-old juvenile or six-week-old adult transgenic plants grown under specified growth conditions were used. Histochemical assay for *β*-*Glucuronidase (GUS)* was performed in the seedlings (whole seedlings, different tissues such as hypocotyl, cotyledon and roots), juvenile and adult plants (rosette leaf and stem). GUS histochemical assay was performed as described (Jefferson et al, 1987) with minor modifications. Approximately 20-25 seedlings were used for histochemical staining. Tissue from the control and transgenic plants were harvested and fixed in fixation buffer (2% formaldehyde, 50 mM sodium phosphate (pH 7.0), 0.05% Triton X-100), and vacuum infiltrated for 4 to 5 min on ice and kept at room temperature for 10 min. The fixation buffer was removed, and the material was washed twice with 50 mM sodium phosphate buffer (pH 7.0) to remove the fixative buffer. The tissue samples were stained using staining buffer [(1.5 mM of X-gluc, 50 mM sodium phosphate (pH 7.0) and 0.1% Triton X-100)] by vacuum infiltrating for 5 to 10 min and then wrapped with aluminium foil and incubated at 37°C overnight in darkness. The next day, samples were observed for staining pattern, and tissue was destained extensively with a destaining solution (Ethanol:Acetic acid: Glycerol at a ratio of 4:4:2) for 10 minutes, and the destained seedlings were stored in 10% glycerol. In a few cases (constant light-to-dark transition experiments), we also used 70% ethanol after overnight staining to remove the chlorophyll for better visualisation of GUS staining. Representative seedlings from different genotypes and growth conditions were aligned on suitable media and photographed.

### GUS spectrometric assay

Approximately 40-50 seedlings from ∼100 mg fresh tissues (seedlings/whole plants/ parts of the plants) were harvested in a micro-centrifuge tube frozen in liquid nitrogen and ground in 100µl of extraction buffer [50 mM sodium phosphate (pH 7.0), 5 mM DTT, 1 mM EDTA, 0.1% sarcosyl, 0.1% Triton X100] at 4 °C. The sample was transferred into a fresh microcentrifuge tube and centrifuged at 10,000 rpm for 5 min at 4 °C. Supernatant was separated and moved to the fresh tube. 10 µl of protein extract was added to the 190 µl of assay buffer (1 mM MUG in extraction buffer) and incubated at 37 °C for 15 min. Next, 180 µl of 0.2 N Na_2_CO_3_ stop buffer was added to a 20 µl reaction mix to stop the reaction. GUS activity was determined by the fluorometric assay as described (Jefferson et al., 1987). Total protein was quantified in the extract using Bradford assay. *β*-*Glucuronidase (GUS)* specific activity was recorded as nanomoles of 4-MU formed per milligram of protein per hour (nanomoles of 4-MU/mg protein/h) formed per milligram of from the initial velocity of the reaction (Jefferson et al., 1987). Finally, the GUS activity was calculated by comparing the spectrophotometric reading to the MU standard and normalizing it to the total protein content and dilution factor wherever applicable.

### Sample collection for tissue-specific GUS activity measurement

Tissue-specific GUS spectrometric assay was carried out from six-day-old seedlings grown in square plates. Hypocotyl, cotyledon and roots were excised separately using a scalpel blade and collected into separate Eppendorf tubes before being frozen in liquid nitrogen. Total protein was extracted, and a GUS spectrometric assay was carried out.

### Luciferase activity assay

We detected luciferase (LUC) activity using Firefly Luciferase Assay Kit 2.0 (Biotium). Approximately 25-30 fresh seedlings (one-week-old) were harvested in a microcentrifuge tube and snap chilled in liquid nitrogen and ground in 150 µl of extraction buffer [50 mM Tris HCl pH 8.0, 150 mM NaCl, 10% glycerol, 5mM DTT, 1% (v/v) Protease Inhibitor Cocktail, 1% NP40, 0.5mM PMSF]. The lysate was cleared by centrifugation at 10,000 rpm for 10 min at 4 °C and transferred into a new tube. The tubes were placed at 4 °C until ready to assay. Firefly working solution was prepared by adding D-Luciferin (10 mg/ml) to assay buffer at a ratio of 1:50. 20 µl of total protein extract was directly added to the plate (Cat. No 781665, Brand 96 well plate). The 100 µl of firefly working solution was added to the protein sample and mixed by gentle pipetting. Then, the plate was immediately placed in a plate reader, and the firefly luminescence measurement was recorded and represented as the average of counts/s^−1^ (cps) per well. Total protein was quantified in the extract using Bradford assay, and the level of luciferase was calculated by normalizing it to the total protein content. The dark condition was maintained during the experiment.

### RNA extraction and gene expression analysis by RT-qPCR

For gene expression analysis using quantitative-PCR (qPCR), RNA was extracted using RNeasy Plant mini kit (QIAGEN) with on-column DNase I digestion according to the manufacturer’s instructions. RNA was quantified using NanoDrop, and approximately 2.0 µg of total RNA was converted into cDNA using a Verso cDNA synthesis kit (Thermo-Fisher scientific) and oligo dT according to the manufacturer’s instructions. Exactly 2.0 µL of 1:20 diluted cDNA was used for qPCR using a 2x SYBR Green Master Mix kit. qPCR experiments were performed in QuantStudio™ 5 Real-Time PCR System. *EF1α* (AT5G60390) was used as an internal control for normalization. Details of the oligonucleotide primers used are provided in **Table S4**.

### Protein extraction, SDS PAGE running and western blot analysis

Approximately 100 mg of tissue was harvested in a microcentrifuge tube, snap frozen in liquid nitrogen and ground in 200 µl of protein extraction buffer [50 mM Tris HCl pH 8.0, 150 mM NaCl, 10% glycerol, 5mM DTT, 1% (v/v) Protease Inhibitor Cocktail, 1% NP40, 0.5mM PMSF]. The protein extract was centrifuged at 10,000 rpm for 15 min at 4°C to pellet down the debris. The supernatant was then transferred to a fresh tube and maintained ice-cold conditions, and an aliquot of 3-5 µl was taken out in a separate tube to estimate protein concentration by Bradford assay. The protein samples diluted to 1µg/µl were boiled for 10 min at 70°C and 50 µg denatured protein samples were loaded onto the SDS PAGE (10% gel)for separating proteins and run for approximately 4 hr. Separated proteins were transferred to the PVDF membrane at 90 volts for 1 hr in transfer buffer (Tris 48 mM, Glycine 39 mM, 20% methanol pH 9.2) in the wet transfer method in the cold room. The membrane was stained with Ponceau-S to confirm the protein transfer and then washed with sterile MQ water. The membrane was then incubated on a rotary shaker for 2 hr in 10 ml blocking buffer (5% non-fat dry milk in Tris-buffered saline (TBS) with 0.05% Tween-20) at room temperature. The blocking reagent was removed, and the affinity-purified primary antibody diluted (1:2500 to 1:10,000) in 10 ml TBS with 0.05% Tween-20 was added and incubated overnight with shaking in the cold room. The next day, the membrane was washed thrice with 10 ml of wash buffer (TBS and 0.05% Tween-20) for 5 min each. The secondary antibody conjugated with HRP diluted (1:5,000 to 10,000) in 10 ml TBS with 0.05% Tween-20 was added and incubated for 2 hr with shaking at room temperature. The membrane was washed thrice with 10 ml of wash buffer at room temperature. The blot was developed using the Super Signal West Femto chemiluminescent substrate kit (Pierce) and following the instructions provided by the manufacturer. A substrate working solution was prepared by mixing peroxide and Luminol/enhancer solutions in a 1:1 ratio. The blot was incubated in that working solution for 5 min in the dark. The blot was removed from the working solution and observed in Chemi-Doc by being exposed for different time durations depending on signal strength. The bands were quantified by ImageJ software. The total PIF4 protein (PIF4 endogenous + PIF4-FLAG) is quantified by considering two bands and normalized to WT (by setting WT to 1). The PIF4-FLAG band was also quantified separately by normalizing to *PIF4-OE1* (by setting *PIF4-OE1* to 1). Actin was used as a loading control, and *pif4-101* was used as a negative control in all the immunoblot experiments. For all the western blots, an identical condition was maintained. The antibody used for detecting endogenous PIF4 level was PIF4 (goat antibody, Agrisera, Cat no. AS163955 at dilution 1:2500). The Actin antibody (abcam, Cat no. ab197345 at dilution 1:10000) was used as the loading control.

### Yeast-one hybrid assay

To generate constructs for yeast one-hybrid assays, bait and prey were prepared. CDS of *PIF4* (encoding full-length protein) from Col-0 cDNA and full-length promoter of *PIF4* (containing G/E-boxes and PBE-box) from genomic DNA of Col-0 were amplified. CDS of *PIF4* was cloned into the pGADT7 vector. The PCR fragment of the promoter of *PIF4* was cloned into *pAbAi* vector, respectively, containing Hind III + Kpn I restriction cut site by restriction-based protocol. The positive colonies were confirmed by colony PCR with gene-specific primers and sequencing by using vector-specific primers. The constructs were transformed into yeast strains *YM4271* and *Yα1867,* and yeast-one hybrid screening was performed in SD/-Ura/-Leu containing 150 ng/ml to 200 ng/m/ Aureobasicidin A for around seven days at 30°C.

### Chromatin Immunoprecipitation (ChIP) assay

ChIP was carried out as described(Kumar et al., 2012) with minor modifications. For this experiment, *PIF4-OE2* seedlings were grown on MS medium for seven days; they were maintained at 22°C under short days (8 h light/16 h dark) in white light. Seedlings (approximately 2.5 gm) were harvested in dim light and directly cross-linked with 1% formaldehyde. ChIP was done using paramagnetic Dynabeads coated with monoclonal anti-FLAG-HRP conjugated antibody (Abcam) following the manufacturer’s instructions. Beads were washed four times with the immunoprecipitation buffer and two washes with Tris-EDTA buffer (TE). Reverse cross-linking was done by boiling at 95°C for 10 min in the presence of 10% Chelex (BioRad) followed by proteinase K treatment at 50°C. The qPCR was performed using SYBR Green dye in QuantStudio™5 Real-Time PCR System (ABI), and enrichment was calculated relative to wild-type controls. PIF4 binding to its promoter was performed using a set of primers spanning the promoter regions covering either PBE-box, E-box or G-box elements. Oligonucleotide sequence details are provided in **Table S4**.

### Sampling of biological samples for various assays

As indicated in respective sections, all experiments were performed with at least two biological replicates. Unless otherwise mentioned, at least 20 seedlings were used for measuring hypocotyl length. ZT23 for SD and ZT4 for LD were followed for harvesting tissue samples for various experiments unless otherwise specified. For the time-course experiment, the seedlings were grown for five days under SD. Tissue was harvested from the sixth day at ZT0 till ZT24 for various durations under both 22°C and 27°C. GUS activity was quantified, and immunoblot experiments were performed. For GUS staining, ∼20-25 seedlings were harvested and stained. The stained representative seedlings were used for the reference picture. For GUS activity measurement, tissue was harvested in two biological replicates, and three technical replicates were used. For gene expression analysis, three independent biological replicates were sampled at specific time points. For qPCR analysis, three technical replicates were used for each biological replicate.

### Mathematical model

Our model is adapted from a published model(Nieto et al., 2020) that we extended by introducing the proposed autoinhibition by PIF4. In our model, *B*(*t*) denotes the photoactivated form of phyB; *E*(*t*) and *C*(*t*) denote the ELF3 and COP1, respectively; and *P*(*t*) denotes the concentration of PIF4. Also, *B*(*t*), *E*(*t*), *C*(*t*), and *P*(*t*) represent the cellular concentrations of respective proteins at a time, *t*. Moreover, *G*(*t*) denotes the hypocotyl length. Here, *G*(*t*) is a coarse-grained variable that represents the collective effect of all growth-promoting genes targeted by PIF4. Additionally, *F*(*t*) denotes the GUS concentration, representing the transgenic promoter activity (i.e., the GUS activity). Note that all molecular concentrations are expressed in arbitrary units, while the hypocotyl length is in *mm*. The following set of ordinary differential equations (ODEs) describe the temporal dynamics of the variables.

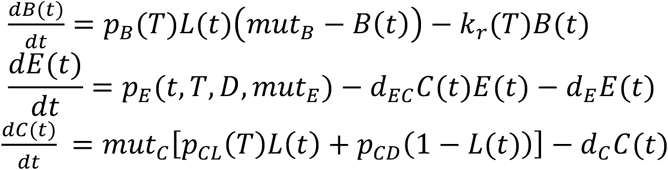

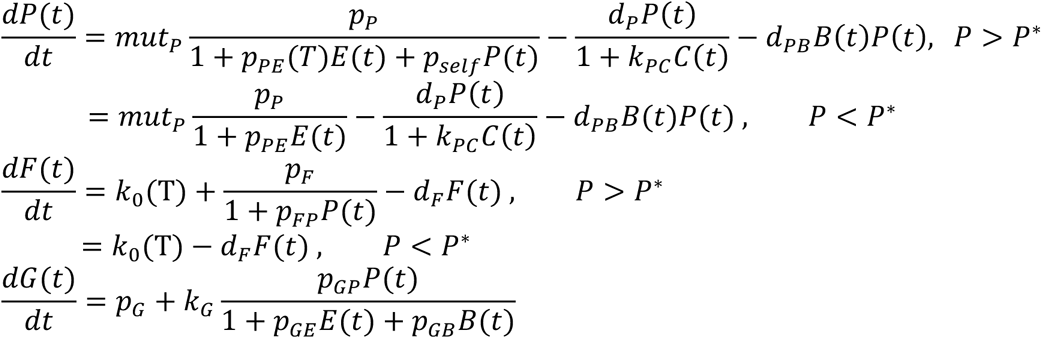

Here, *L*(*t*) is a binary variable denoting the presence or absence of light, which is either 0 in dark or 1 in white/red light. In the ODEs (Equation 1), decay rates are, in general, denoted by *d_K_* where *K* symbolically represents the variable *K* (see the detailed parameter descriptions in **Table S1** and **S2**). Similarly, *p_K_*, in general, denotes the production rate of the variable *K*. Specifically, the production rate of ELF3 (denoted by *p_E_*) is modelled by an oscillatory function (see Equation 2). The production rates of COP1 are denoted by *p_CL_* and *p_CD_*, in light and dark, respectively. Notably, we included a multiplicative factor *mut_K_* that modifies the production rate of variable *K* in the corresponding knock-out and over-expressor lines, i.e., *mut_K_* = 1 for the wild type, *mut_K_* < 1 for the mutants, and *mut_K_* > 1 for overexpresson lines (see **Table S2** for specific values of these multipliers in different genotypes). In general, the parameter, *p_MN_*, denotes the interaction strength between variables *M* and *N* (see the detailed descriptions in **Table S1** and **S2**).

Moreover, we assumed that PIF4 autoinhibition occurs above a threshold PIF4 concentration denoted by *P*^∗^. Consequently, PIF4-induced inhibition of GUS also takes place when PIF4 concentration is above *P*^∗^. Note that the threshold PIF4 concentration (*P*^∗^) depends on a specific genotype, in general. The intensity or strength of autoinhibition for PIF4 synthesis is denoted by *p_self_*. Furthermore, *k*_0_ is the basal production rates of GUS. To model the hypocotyl growth, we assumed that *k_G_* is a coarse-grained parameter representing the conversion of PIF4 target gene expressions into the overall growth. Finally, some parameters are assumed to depend on the temperature, *T*, and have different values at 22°C and 27°C (see **Table S1** and **S2**).

Previous studies(Nieto et al., 2020) showed that ELF3 has an oscillatory nature depending on the diurnal days. Thus, we incorporated the ELF3 synthesis by an oscillatory function as below:

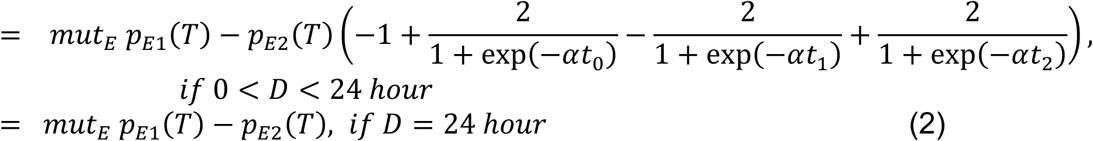

Here, D represents the time of light exposure (i.e., the day length). The production rate of ELF3 oscillates between (*p_E_*_1_ + *p_E_*_1_) and (*p_E_*_1_ − *p_E_*_1_). Also, *t*_0_ = *mod* (t, 24), *t*_1_ = *t*_0_ − *D*, *t*_2_ = *t*_0_ − 24. The parameter *α* defines the sharpness of transition between the maximum and minimum value of *p_E_*.

### Simulation details

We numerically solved the ODEs (Equation 1) using MATLAB (ode45 solver). Initially, all the variables were kept at zero for the numerical solution. We took some model parameters from a previous study (Nieto et al., 2020), while other parameters were obtained by fitting to the data of hypocotyl lengths from wildtype seedlings grown under 4, 8, 12, and 16-hours white light cycles at 22°C (see the details of parameter selection and fitting procedure below). Then, we predicted the hypocotyl lengths and GUS activities for other genotypes by varying the parameters, *mut_K_*, *P*^∗^, and *p_self_* (as in **Table S2**).

### Parameter selection and fitting

We first fixed some of the parameters from a previous study (Nieto et al., 2020), on which our model is based (see **Table S1**). Next, there were 7 more unknown parameters, which were estimated from the ‘non-linear least squares’ fitting to the wild-type (WT) hypocotyl data at 22^7^C temperature. Also, note that all multiplicative factors for the WT are unity by definition (*mut_K_* = 1).

The non-linear least squares method was implemented using the open-source Python package *scipy.optimize.curve_fit* (Version SciPy v1.12.0). To implement this method, the number of data points should be more or at least equal to the number of unknown parameters. However, we have only four experimental data points of mean hypocotyl lengths (each averaged over ∼20 replicates) and seven unknown parameters. Due to the limited number of experimental data points, we have considered all ∼ 20 replicates to estimate the parameter uncertainty (given by standard deviation in **Table S2**), and the corresponding goodness of fit (given by *R*^2^ values). The *R*^2^ values for fitting the model with experimental data points are shown in Figure 2B.

After determining all parameters for the WT, we did not alter some of the parameters for other genotypes, namely, inhibition rate of ELF3 by COP1 (*d_EC_*), basal GUS production rate (*k*_0_), GUS production rate influenced by PIF4 (*p_F_*), GUS decay rate(*d_F_*), and intensity of PIF4’s inhibition of GUS (*p_FP_*). For all mutants and overexpression lines, we varied only three parameters, PIF4 threshold concentration for autoinhibition (*P*^∗^),negative feedback strength (*p_self_*), and the multiplicative factors (*mut_K_*), to again estimate their values by fitting to the corresponding hypocotyl data for each genotype (see **Table S2**).

Finally, note that all parameter values (with corresponding standard deviation) were obtained by fitting only for 22°C temperatures. However, the values of threshold (*P*^∗^) and feedback strength (*p_self_*) were assumed to change slightly at 27°C temperature (without any fitting).

### Quantification and statistical analysis

The data shown are the mean±SD. The number of biological replicates (n) and the statistical details of each experiment are indicated in the corresponding figure legend. For the hypocotyl length measurement and quantification of western blots, Image J software was used. GraphPad Prism 8.0 and Microsoft Excel were used for graph preparation and statistical analysis. The statistical significance between or among treatments and/or genotypes was determined based on one-way or two-way ANOVA followed by Tukey’s HSD test (P<0.05). Significant differences between genotypes and/or temperatures are denoted by different letters. For immunoblot data, all the experiments were repeated three times, and the data from the representative experiment is shown.

## Acknowledgements

This work is supported by grants from the Department of Biotechnology (Ramalingaswami Re-entry Fellowship grant, BT/RLF/Re-entry/ 28/2017), Science and Engineering Research Board (start-up research grant, SRG/2019/000446), IISER Kolkata Intramural grant (Ministry of Education, Government of India) to S.N.G. We thank Dr. Vinod Kumar for *pPIF4:PIF4-FLAG* and Prof. Salome Prat for the *pPIF4:LUC* seeds.. The graphical abstract is created with Biorender (https://app.biorender.com/illustrations). We also thank Sumana Annagiri’s lab (IISER Kolkata) for the microscopy facility. S.D. acknowledges IISER Kolkata, and V.G. and K. M. acknowledge the University Grants Commission (UGC, Govt of India) for their doctoral fellowship.

## Author contributions

S.N.G. conceived and supervised the project, designed experiments, and analyzed the data. S.D. and V.G. designed and performed the experiments, analysed the data, prepared figures and wrote the first draft of the manuscript. K.M. and D.D. developed the mathematical model. S.N.G supervised experiments, and D.D supervised theoretical modelling. S.N.G. and D.D. wrote the final manuscript with help from S.D., V.G., and K.M. All the authors read and approved the final draft.

## Declaration of interests

The authors declare no competing interests.

## Supporting Information

**Figure S1.**
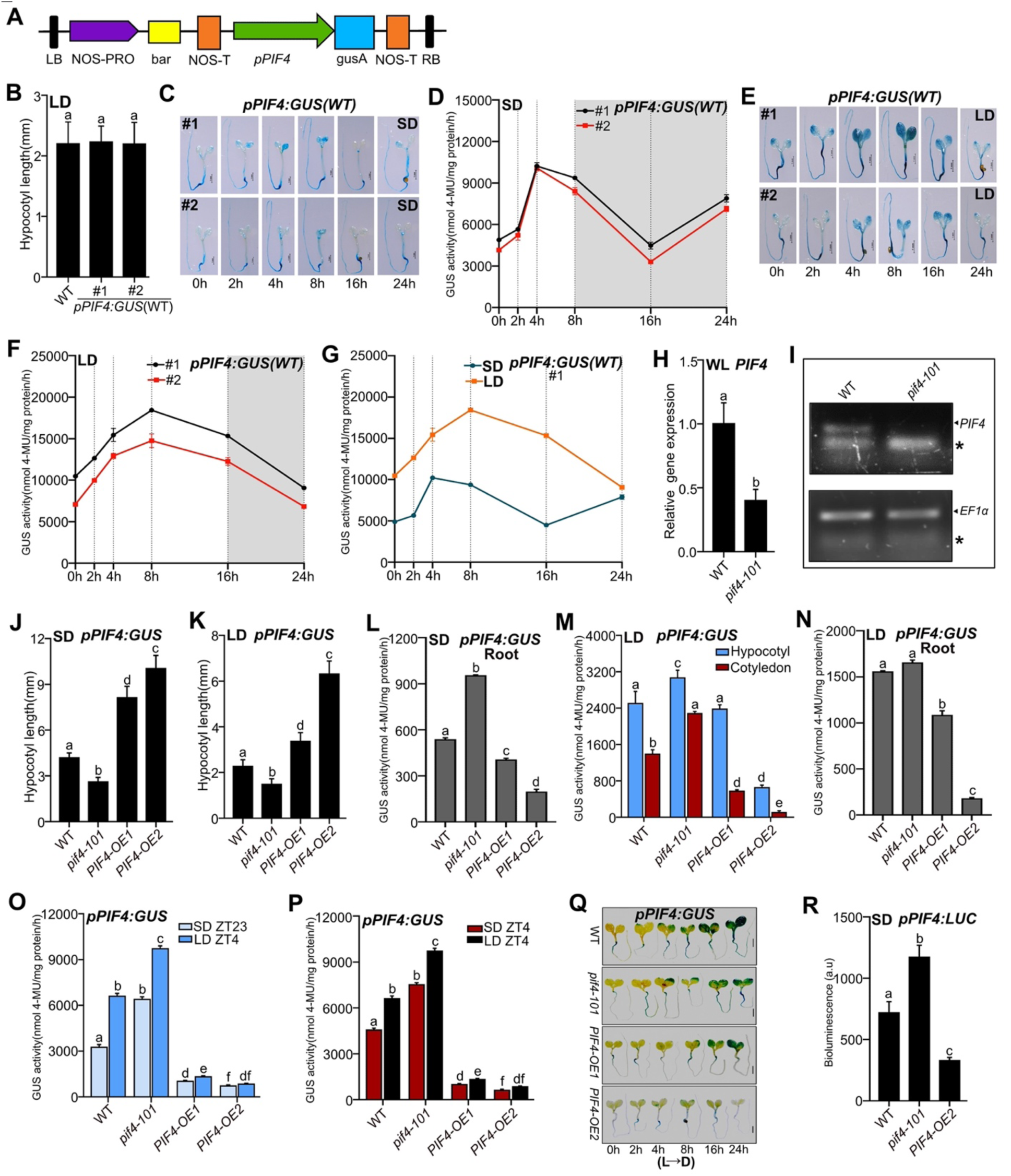
PIF4 negatively autoregulates its own gene expression in a photoperiod-dependent manner. (A) Schematic diagram of the construct used for making *pPIF4:GUS*. (B) Two independent *pPIF4:GU*S lines under WT background are used. Hypocotyl length of six-day-old WT, *pPIF4:GU*S line #1 and #2 grown in LD under 22°C. (C-F) GUS stained images and GUS activities from *pPIF4:GUS* line #1 and #2 grown under SD (C-D) and LD (E-F) conditions, respectively. Tissue was harvested at ZT0 on the sixth day and grown for 24 hours. The dark period is shown in grey in the graph. (G) Measurement of GUS activity of *pPIF4:GUS* line #1 grown under SD and LD in diurnal conditions. (H) PIF4 transcript level was measured by qPCR in six-day-old WT, and *pif4-101* mutants were grown in SD at ZT23. The transcript level was normalized to WT. (I) Semi-quantitative PCR analysis of PIF4 for WT and *pif4-101* mutant. The diluted cDNA was run in 2% agarose gel. The asterisk denotes a non-specific band. EF1α was used as a loading control. (J and K) Hypocotyl length of WT, *pif4-101*, *PIF4-OE1* and *PIF4-OE2* carrying the transgene *pPIF4:GUS* seedlings grown in SD (J) and LD (K). (L) Root was dissected from ten-day-old whole seedlings of indicated genotypes grown in vertical plates under SD conditions (ZT23), and quantitative GUS activity was measured from the roots. (M and N) The GUS activity in hypocotyl, cotyledon (M) and root (N) harvested from six-day-old seedlings under LD (tissue was harvested at ZT4). (O) GUS activity was calculated from six-day-old seedlings grown under SD and LD. Tissue was harvested at the end of the night for SD (ZT23) and daytime for LD (ZT4). (P) Quantification of GUS activity from six-day-old seedlings grown under SD and LD in WL (ZT4). (Q) GUS-stained images of constant light (22°C) grown seedlings shifted to dark for indicated genotypes at various intervals. (R) Microplate bioluminescence detection of WT, *pif4-101* and *PIF4-OE2* lines expressing the *pPIF4:LUC*. The six-day-old seedlings were grown in SD photoperiod at 22°C. Seedlings were harvested at the end of the night (ZT23). Values represent mean ± SD of the 1-s absolute bioluminescence of at least 24 seedlings per genotype. The unit of bioluminescence was represented as an arbitrary unit (a.u.). Related to Figure 1.

**Figure S2.**
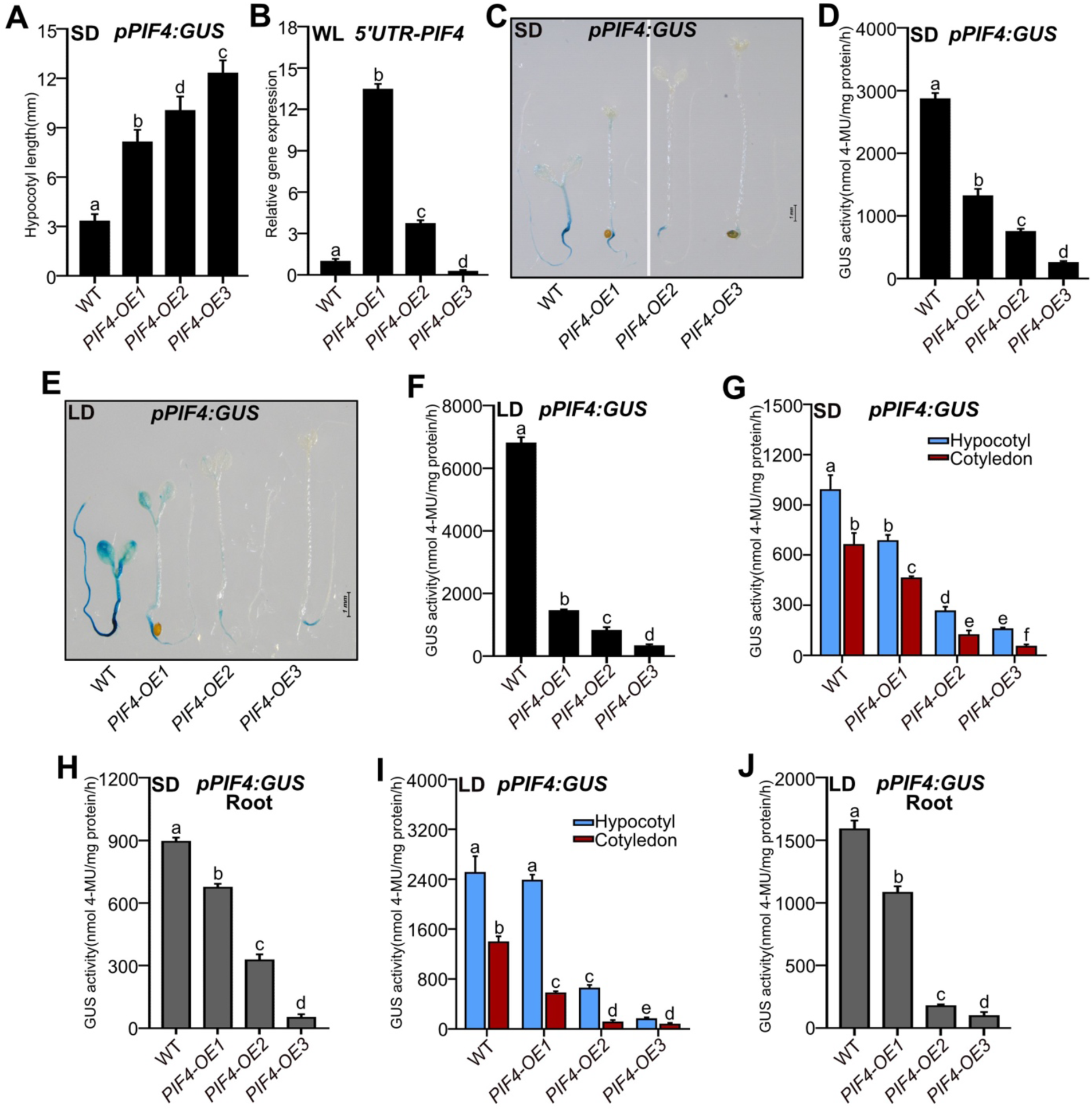
Constitutive overexpression of PIF4 results in stronger autoinhibition of PIF4 promoter activity. (A) Hypocotyl length of six-day-old WT, *PIF4-OE1, PIF4-OE2* and *PIF4-OE3* seedlings carrying the transgene *pPIF4:GUS* grown in SD. (B) Relative *PIF4* transcript levels (at ZT23) as revealed by qRT-PCR using primer set (5’ UTR+ exon) specific to endogenous PIF4 gene in six-day-old WT, *PIF4-OE1, PIF4-OE2* and *PIF4-OE3* under SD in WL. *EF1α* was used as a loading control. The transcript level of each sample was normalized to WT. Error bars depict the mean ± SD of three biological replicates. (C-F) GUS staining and activity measurement of six-day-old seedlings grown under SD at ZT23 (C and D) and LD at ZT4 (E and F) conditions. The vertical line in C indicates two separate images. (G-J) GUS activities were quantified from different tissues from six-day-old seedlings grown under SD (G and H) and LD (I and J). The seedlings were separated into hypocotyl, cotyledon and root, and the GUS activity was measured separately. All bar graphs represent data (mean ± SD) from three biological replicates. Different letters indicate a significant difference (one-way ANOVA and two-way ANOVA with Tukey’s HSD test, P < 0.05, n ≥ 40 seedlings for GUS activity data). Related to to Figure 1.

**Figure S3.**
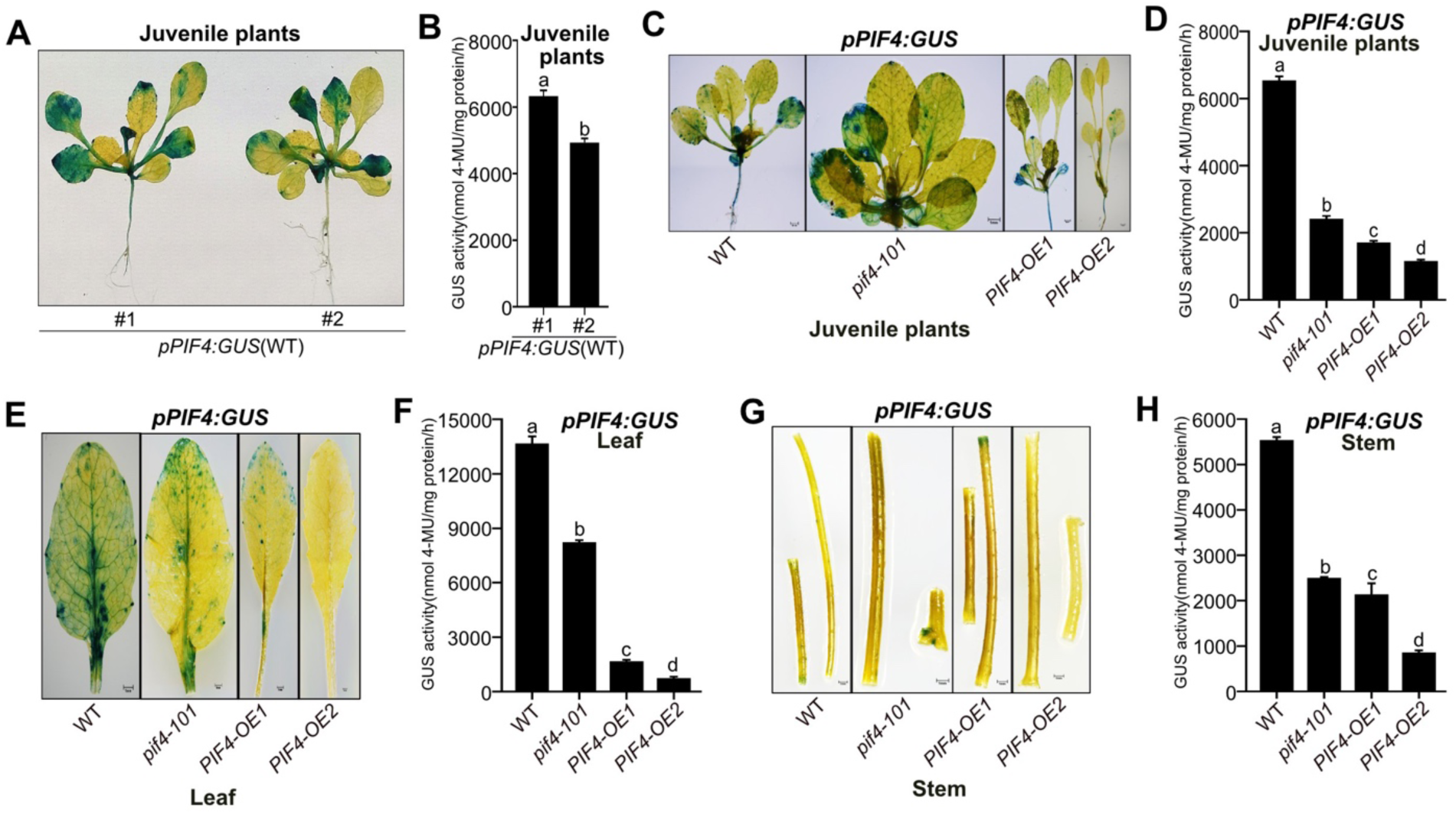
Auto-inhibition of the PIF4 promoter persists in the adult stage. (A and B) GUS stained images (A) and GUS activity (B) of three-week-old juvenile plants of two independent *pPIF4:GUS* lines #1 and #2 under WT background grown in LD conditions. (C-H) Tissue-specific expression patterns. Representative GUS staining and activity in three-week juvenile plants (C and D) and in rosette leaves (E and F) and stem (G and H) of six-week-old adult plants grown at 22°C under LD and tissue was harvested at ZT4. Data represent mean±SD (n=6). Different letters indicate significant differences (one-way ANOVA with Tukey’s HSD test, P < 0.05). Related to Figure 1.

**Figure S4.**
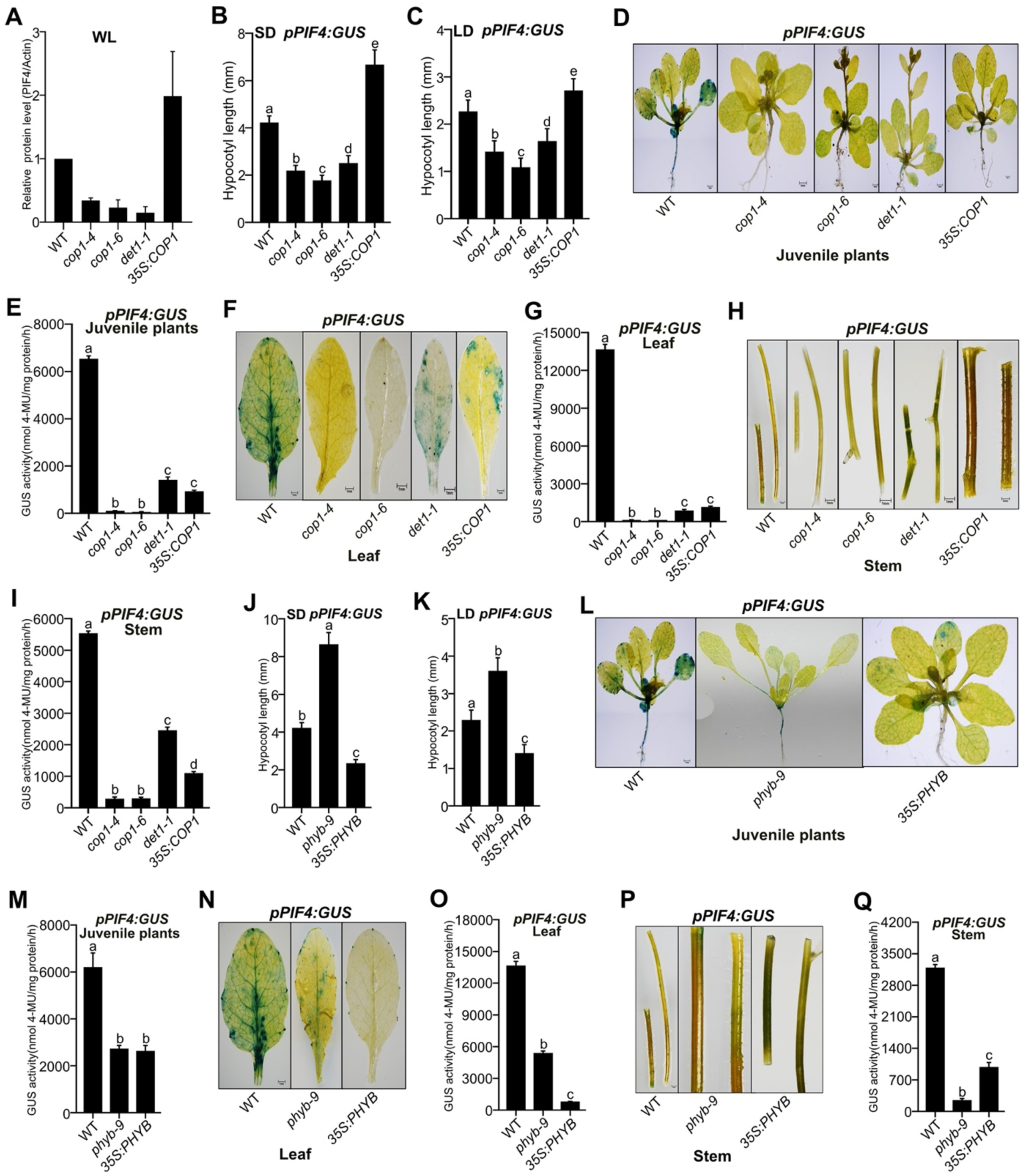
COP1/DET1 promotes, while PHYB inhibits PIF4 autoinhibition. (A and B) Hypocotyl length of six-day-old WT, *cop1-4, cop1-6, det1-1* and *35S:COP1* seedlings carrying the transgene *pPIF4:GUS* grown in SD (A) and LD (B). (C-H) GUS staining and GUS activity from the above-mentioned genotypes are detected in three-week-old juvenile plants (C and D), rosette leaves (E and F) and stem (G and H) of six-week-old adult plants grown at 22°C under LD. Whole plant tissue was harvested for staining and activity at ZT4. (I and J) Hypocotyl length of six-day-old WT, *phyb-9* and *35S:PHYB* seedlings carrying the transgene *pPIF4:GUS* grown in SD (I) and LD (J). (K-P) Representative GUS staining and GUS activity from three-week-old juvenile plants (K and L) and rosette leaves (M and N), stem (O and P) of six-week-old adult plants grown at 22°C under LD. Whole plant tissue was harvested at ZT4. Data represent mean±SD (n>20 for Hypocotyl length experiment and n=6 for adult plant histochemical assay). Different letters indicate significant differences (one-way ANOVA with Tukey’s HSD test, P < 0.05). Related to Figures 3 and 4.

**Figure S5.**
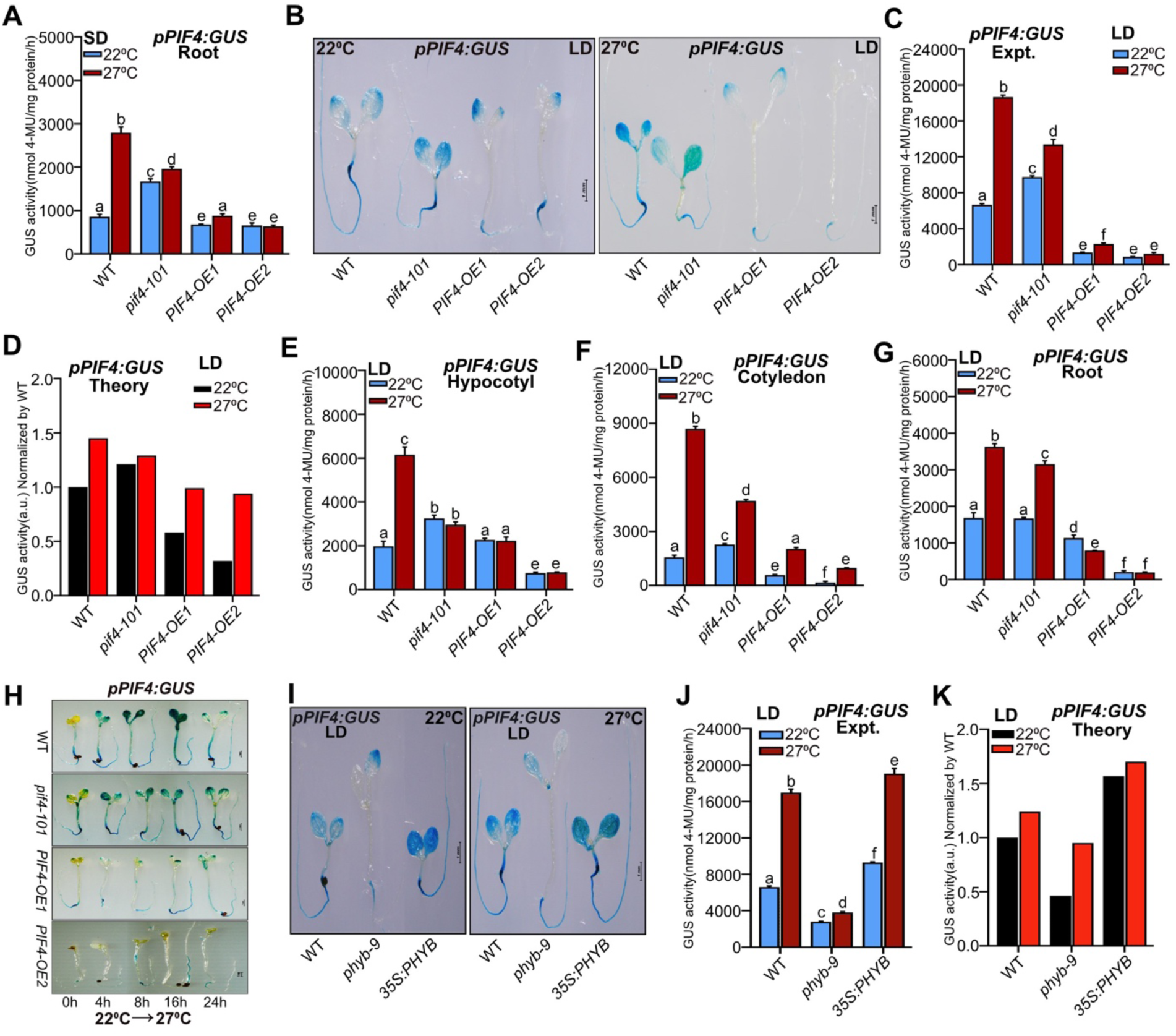
Temperature-mediated autoinhibition of PIF4 promoter activity is dependent on endogenous PIF4 protein concentration. (A) GUS activities from the dissected root from six-day-old whole seedlings grown under SD at 22°C and 27°C. (B-D) Representative GUS-stained image (B), experimental GUS activity (C), and model-predicted GUS activity (D) of six-day-old seedlings grown under LD in WL at 22°C and 27°C. (E-G) GUS activities measured from hypocotyl (E), cotyledon (F) and roots (G) detached from whole seedlings grown under 22°C and 27°C in LD. (H) Representative GUS-stained images of WT, *PIF4-OE1* and *PIF4-OE2* grown under 22°C and shifted to 27°C for the mentioned time point under SD. (I-K) Representative GUS-stained images of WT, *phyb9,* and *35S:PHYB-OE* carrying the transgene *pPIF4:GUS* are shown (I), experimental GUS activity (J) and model predicted GUS activity (K) measurement under 22°C and 27°C in LD at WL. (K) Representative GUS-stained images of WT, *PIF4-OE1* and *PIF4-OE2* grown under 22°C and shifted to 27°C for the mentioned time point under SD. For GUS staining and activity measurement, the samples were harvested at ZT23 for SD and ZT4 for LD conditions. Data represent mean±SD. Different letters indicate significant differences (two-way ANOVA with Tukey’s HSD test, P < 0.05). Related to Figure 5.

**Table S1.** The description of parameters and corresponding values. The values are taken from (Nieto et al., 2020), and they are fixed for all genotypes. Note that the shaded boxes denote the parameters that slightly change from 22°C to 27°C.

| Description of parameters | Values at 22°C | Values at 27°C |
| --- | --- | --- |
| Light-induced activation rate of phyB | $p_B = 10.0$ | $p_B = 0.860$ |
| Deactivation rate of phyB in dark | $k_r = 0.232$ | $k_r = 0.411$ |
| Parameter related to the production rate of ELF3 (see Equation 2) | $p_{E1} = 108$ | $p_{E1} = 127$ |
| Parameter related to the production rate of ELF3 (see Equation 2) | $p_{E2} = 39.8$ | $p_{E2} = 7.29$ |
| Production rate of COP1 in light | $p_{CL} = 1.00$ | $p_{CL} = 5.37$ |
| Intensity of ELF3's inhibition of PIF4 production | $p_{PE} = 0.332$ | $p_{PE} = 0.028$ |
| Production rate of COP1 in dark | $p_{CD} = 112$ | $p_{CD} = 112$ |
| Decay rate of ELF3 | $d_E = 27.2$ | $d_E = 27.2$ |
| Production rate of hypocotyl growth | $p_{GP} = 2.93$ | $p_{GP} = 2.93$ |
| Decay rate of COP1 | $d_C = 1.79$ | $d_C = 1.79$ |
| Decay rate of PIF4 | $d_P = 4.91$ | $d_P = 4.91$ |
| Basal rate of hypocotyl growth | $p_G = 0.009$ | $p_G = 0.009$ |
| Intensity of phyB's inhibition of hypocotyl growth | $p_{GB} = 10.7$ | $p_{GB} = 10.7$ |
| Inhibition rate of PIF4 by phyB | $d_{PB} = 0.313$ | $d_{PB} = 0.313$ |
| Conversion factor between PIF4-targeted gene expression and hypocotyl growth | $k_G = 0.113$ | $k_G = 0.113$ |
| Intensity of ELF3's inhibition of hypocotyl growth | $p_{GE} = 0.465$ | $p_{GE} = 0.465$ |
| Intensity of COP1's inhibition of PIF4 degradation | $k_{PC} = 34.3$ | $k_{PC} = 34.3$ |
| Parameter defining the sharpness of transitions between maximum and minimum production rates of ELF3 | $\alpha = 5$ | $\alpha = 5$ |
| Production rate of PIF4 | $p_P = 1$ | $p_P = 1$ |

**Table S2:** Parameter values estimated (with uncertainty) from non-linear least squares fitting. Note that the shaded boxes represent the parameter values obtained by fitting to the wildtype data and kept unchanged for all other genotypes.

| Genotypes | Description of parameters | Values at 22°C<br>(Estimated) | Values<br>at 27°C<br>(Assumed) |
| --- | --- | --- | --- |
| **WT** | Inhibition rate of ELF3 by COP1 | $d_{EC} = 0.01 \pm 0.0133$ | $d_{EC} = 0.01$ |
| | Basal rate of GUS production | $k_0 = 10 \pm 0.2007$ | $k_0 = 10$ |
| | Production rate of GUS | $p_F = 450 \pm 10.7729$ | $p_F = 450$ |
| | Decay rate of GUS | $d_F = 0.0009 \pm 0.0206$ | $d_F = 0.0009$ |
| | Intensity of PIF4's inhibition of GUS expression | $p_{FP} = 22 \pm 0.03975$ | $p_{FP} = 22$ |
| | Multiplier<br>($mut_k = 1; k = E, B, P, C$) | $mut_k = 1$ | $mut_k = 1$ |
| | PIF4 threshold concentration | $P^* = 0.2 \pm 0.00001$ | $P^* = 0.2$ |
| | Negative feedback strength | $p_{self} = 25 \pm 0.3534$ | $p_{self} = 20$ |
| ***pif4-101*** | Multiplier | $mut_P = 0.65 \pm 0.0214$ | $mut_P = 0.65$ |
| | PIF4 threshold concentration | $P^* = 0.04 \pm 0.0242$ | $P^* = 0.04$ |
| | Negative feedback strength | $p_{self} = 25 \pm 0.5818$ | $p_{self} = 5$ |
| ***PIF4-OE1*** | Multiplier | $mut_P = 4 \pm 1.0599E - 09$ | $mut_P = 4$ |
| | PIF4 threshold concentration | $P^* = 0.2 \pm 1.58E - 09$ | $P^* = 0.2$ |
| | Negative feedback strength | $p_{self} = 25 \pm 0.00031$ | $p_{self} = 20$ |
| ***PIF4-OE2*** | Multiplier | $mut_P = 5.9 \pm 0.0000001$ | $mut_P = 5.9$ |
| | PIF4 threshold concentration | $P^* = 1.3 \pm 3.88E - 08$ | $P^* = 1.3$ |
| | Negative feedback strength | $p_{self} = 25 \pm 4.350E - 07$ | $p_{self} = 20$ |
| ***phyb-9*** | Multiplier | $mut_B = 0.6 \pm 0.000021$ | $mut_B = 0.6$ |
| | PIF4 threshold concentration | $P^* = 0.6 \pm 2.07E - 09$ | $P^* = 0.6$ |
| | Negative feedback strength | $p_{self} = 25 \pm 3.021E - 07$ | $p_{self} = 5$ |
| ***35S:PHYB*** | Multiplier | $mut_B = 4 \pm 0.0585$ | $mut_B = 4$ |
| | PIF4 threshold concentration | $P^* = 0.1 \pm 0.0136$ | $P^* = 0.1$ |
| | Negative feedback strength | $p_{self} = 25 \pm 0.5215$ | $p_{self} = 23$ |
| ***cop1-4*** | Multiplier | $mut_C = 0.001 \pm 0.00006$ | ----- |
| | PIF4 threshold concentration | $P^* = 0.35 \pm 2.213E - 20$ | ----- |
| | Negative feedback strength | $p_{self} = 25 \pm 1.48E - 07$ | ----- |
| ***35S:COP1*** | Multiplier | $mut_C = 50 \pm 0.61114$ | ----- |
| | PIF4 threshold concentration | $P^* = 0.6 \pm 0.01429$ | ----- |
| | Negative feedback strength | $p_{self} = 25 \pm 0.32414$ | ----- |

**Table S3:** Parameter values to produce the dynamics of PIF4 protein and GUS activity (corresponding to Figures 6G and 6H). Shaded boxes represent the unchanged parameter values for all genotypes.

| Genotypes | Description of parameters | Values at 22 Degree | Values at 27 Degree |
| --- | --- | --- | --- |
| **WT** | Light-induced activation rate of phyB | $P_B = 10$ | $P_B = 5$ |
| | Deactivation rate of phyB in dark | $k_r = 0.232$ | $k_r = 0.251$ |
| | Production rate of COP1 in light | $p_{CL} = 1$ | $p_{CL} = 2.37$ |
| | Basal rate of GUS production | $k_0 = 10$ | $k_0 = 100$ |
| | Negative Feedback strength at the beginning of the day | $P_{self}^{max/day} = 70$ | $P_{self}^{max/day} = 20$ |
| | Negative feedback strength in the midday | $P_{self}^{min} = 0.1$ | $P_{self}^{min} = 0.1$ |
| | Negative feedback strength at night | $P_{self}^{max/night} = 80$ | $P_{self}^{max/night} = 20$ |
| | PIF4-mediated GUS inhibition rate at the beginning of the day | $P_{FP}^{max/day} = 30$ | $P_{FP}^{max/day} = 5$ |
| | PIF4-mediated GUS inhibition rate in the midday | $P_{FP}^{min} = 0.5$ | $P_{FP}^{min} = 0$ |
| | PIF4-mediated GUS inhibition rate at night | $P_{FP}^{max/night} = 10$ | $P_{FP}^{max/night} = 5$ |
| **PIF4-OE2** | Basal rate of GUS production | $k_0 = 1$ | $k_0 = 20$ |
| | Negative Feedback strength at the beginning of the day | $P_{self}^{max/day} = 90$ | $P_{self}^{max/day} = 85$ |
| | Negative feedback strength in the midday | $P_{self}^{min} = 1$ | $P_{self}^{min} = 0.95$ |
| | Negative feedback strength at night | $P_{self}^{max/night} = 110$ | $P_{self}^{max/night} = 100s$ |
| | PIF4-mediated GUS inhibition rate at the beginning of the day | $P_{FP}^{max/day} = 300$ | $P_{FP}^{max/day} = 300$ |
| | PIF4-mediated GUS inhibition rate in the midday | $P_{FP}^{min} = 50$ | $P_{FP}^{min} = 30$ |
| | PIF4-mediated GUS inhibition rate at night | $P_{FP}^{max/night} = 300$ | $P_{FP}^{max/night} = 300$ |

**Table S4.**
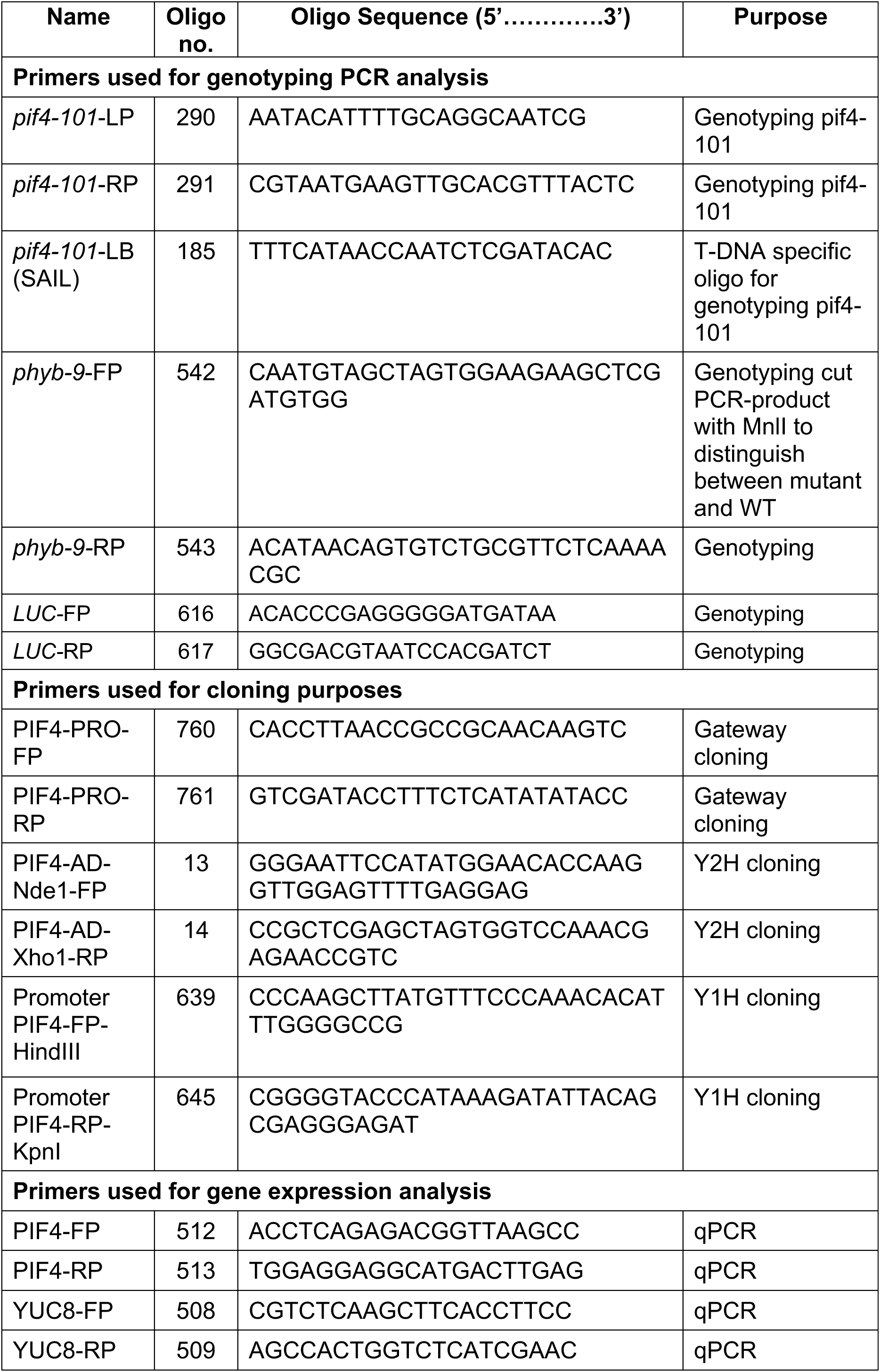
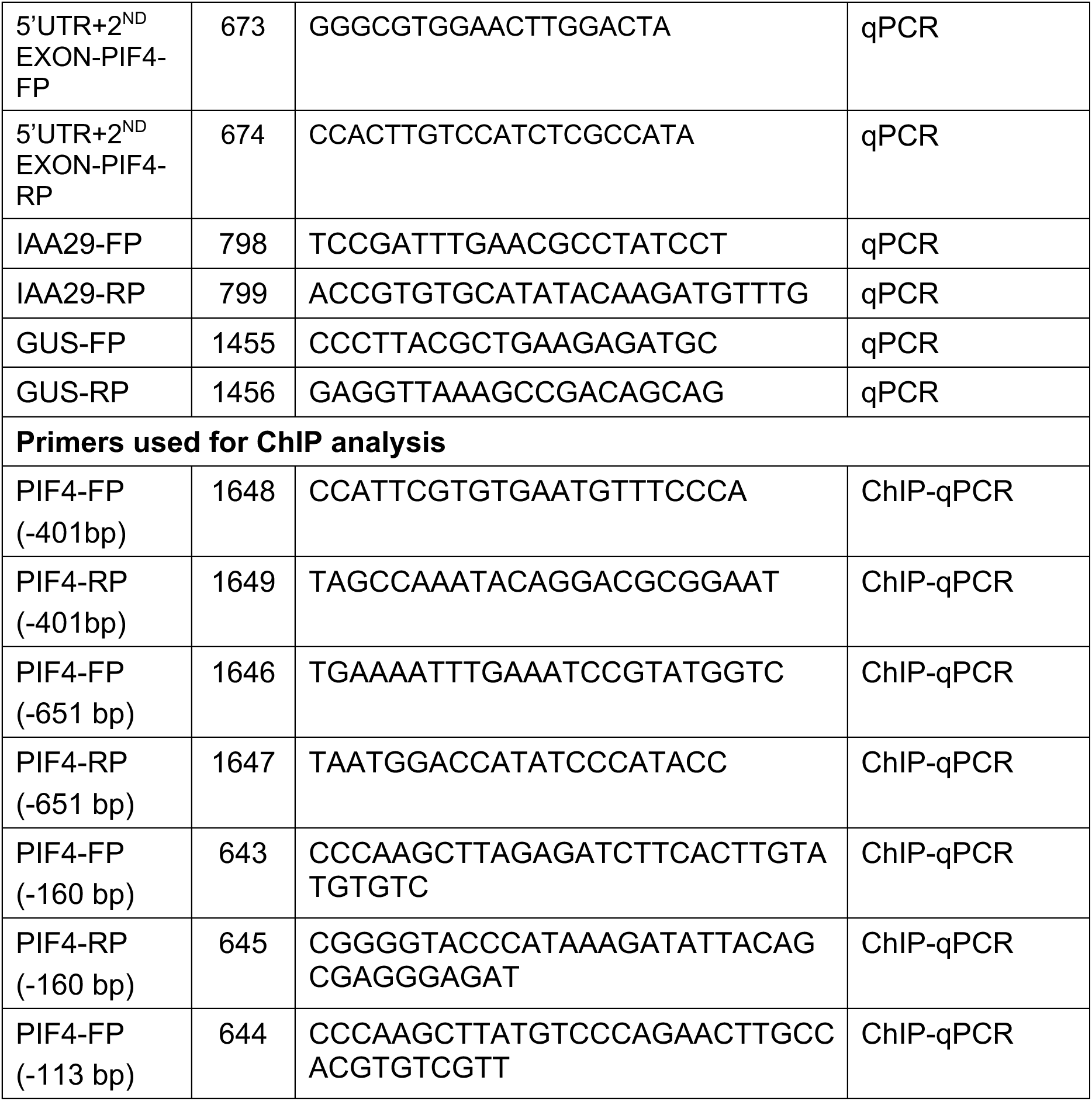
List of oligonucleotide sequences used in the study.

## Notes

### Competing Interest Statement

The authors have declared no competing interest.

